# MSK-mediated phosphorylation of Histone 3 Ser28 couples MAPK signaling with early gene induction and cardiac hypertrophy

**DOI:** 10.1101/2020.07.28.224832

**Authors:** Emma L. Robinson, Faye M Drawnel, Saher Mehdi, Caroline R. Archer, Wei Liu, Hanneke Okkenhaug, Kanar Alkass, Jan Magnus Aronsen, Chandan K Nagaraju, Ivar Sjaastad, Karin R Sipido, Olaf Bergmann, J. Simon C. Arthur, Xin Wang, H. Llewelyn Roderick

## Abstract

Heart failure is a leading cause of death that develops subsequent to deleterious hypertrophic cardiac remodelling. MAPK pathways play a key role in coordinating the induction of gene expression during hypertrophy. Induction of the immediate early gene (IEG) response is a necessary and early event in this process. How MAPK and IEG expression are coupled during cardiac hypertrophy is not yet resolved. Here, in vitro, in rodent models and in human samples, we demonstrate that MAPK-stimulated IEG induction depends on the Mitogen and Stress activated protein Kinase (MSK) and its phosphorylation of histone H3 at serine 28 (pH3S28). pH3S28 in IEG promoters in turn recruits Brg1, a BAF60 ATP-dependent chromatin remodelling complex component, initiating gene expression. Without MSK activity and IEG induction, the hypertrophic response is suppressed. These studies provide new mechanistic insights and highlight the role of signalling to the epigenome in gene expression regulation during cardiac hypertrophy.

**Brief summary one sentence:** MSK1/2 phosphorylation of Histone 3 Serine 28 couples MAPK signalling with chromatin remodelling and immediate early gene expression to induce pro-hypertrophic cardiac transcriptional responses.

## Introduction

Cardiovascular diseases (CVDs) are the leading cause of mortality and morbidity worldwide (Savarese and Lund, 2017). While cardiac hypertrophy is initially an adaptive response to increased workload or stress that enables cardiac function to meet the changing needs of the organism, when induced by pathological cues such as aortic stenosis, prolonged hypertension or myocardial infarction, over time, the response can decompensate resulting in a decline in cardiac function and progression to heart failure. At the cellular level, owing to their terminally differentiated status, hypertrophy of the cardiac muscle is mediated by growth of existing cardiomyocytes and not through their proliferation (Alkass et al., 2015).

Signalling pathways activated downstream of G protein coupled receptors (GPCR), such as those liganded by endothelin-1 (ET-1) and angiotensin II play a fundamental role in the induction of pathological hypertrophic remodelling (Drawnel et al., 2013; Wang et al., 2018). Owing to their nodal position in pathways downstream of GPCR activation, Mitogen Activated Protein Kinases (MAPK) are of particular importance in induction of hypertrophic growth of cardiomyocytes. Indeed, these signalling mediators affect pathological hypertrophic growth via regulation of protein synthesis, cell survival, metabolism and gene transcription. (Bueno et al., 2000; Rose et al., 2010).

MAPKs fall into four major families which are categorised according to the terminal kinase in the pathway – the extracellular regulated kinases 1 and 2 (ERK1/2), p38 MAPK, c-Jun N-terminal kinases 1 and two (JNK1/2) and ERK5 (Bueno et al., 2000; Rose et al., 2010). Involvement of all limbs of this family of kinases has been demonstrated in regulating hypertrophic remodelling, the phenotypic outcome differs depending upon the relative activation of each family member (Garrington and Johnson, 1999; Heineke and Molkentin, 2006; Rose et al., 2010). For example, enhanced ERK signalling leads to concentric cardiac hypertrophy with preserved cardiac function, whereas the alpha isoform of p38 has a pro-apoptotic function, which when prevented, results in protection against CM-associated cardiac injury (Bueno et al., 2000; Kaiser et al., 2004; Marber et al., 2011; Yokota and Wang, 2016).

During cardiac hypertrophy, MAPK pathways intervene in transcriptional regulation via phosphorylation-dependent modulation of transcription factors including NFAT, Elk, SRF and GATA4 (Sanna et al., 2005). Preceding expression of hypertrophic genes, ERK-dependent activation of an immediate early gene (IEG) response is observed in cardiomyocytes. Specifically, IEG expression is rapidly activated in cultured rat neonatal ventricular cardiomyocytes in response to stimulation with hypertrophic agonists, and in vivo following pressure overload (Archer et al., 2017; Iwaki et al., 1990; Izumo et al., 1988). This response is initiated through phosphorylation-dependent activation of the AP-1 family of transcription factors, which promote the induction of IEG expression within minutes of the initiating cue (Balmanno and Cook, 1999; Gille et al., 1992; Karin et al., 1997). Proto-typical IEG include members of the FOS (c-FOS, FOSB, FRA-1 and FRA-2), Jun (JUNB, JUND and c-JUN) and activating transcription factor (ATF; ATFa, ATF2, LRF1/ATF3, ATF4 and B-ATF) families of basic-leucine zipper transcription factors (Eferl and Wagner, 2003). Heterodimers of Fos and Jun associate with ATFs to form the AP-1 transcription factor complex (Glover and Harrison, 1995). The actions of AP-1 transcription factors are also mediated via direct interactions with other transcription factors, many of which, including NFκB and NFAT are involved in collaborations in regulating gene expression in many non-cardiac systems (Chen et al., 1998; Torgerson et al., 1998; Yang et al., 2010). The AP-1 complex is also involved in cardiac hypertrophic responses. This role is clearly illustrated in in vitro and in vivo experiments in which AP-1 activity is suppressed through expression of dominant negative JUN (DN-JUN; also known as TAM67) or of the endogenous AP-1 inhibitor JUND (Hilfiker-Kleiner et al., 2006; Kim-Mitsuyama et al., 2006; Takeuchi et al.). Moreover, muscle-specific knockout of c-Jun results in a loss of the initial compensatory phase of the pathological hypertrophic response and a progression to cardiac dilation, indicating a requirement for c-Jun in the initial phase of the cardiac hypertrophic response (Tachibana et al., 2006; Windak et al., 2013). Taken together, these results indicate that despite certain specific roles for different IEG, the AP-1 transcription factor complex fulfils important and complex functions in hypertrophic remodelling.

Although ERK activation and IEG induction in cardiomyocytes are highly correlated, the mechanism by which these events are coupled during induction of cardiomyocyte hypertrophy is not resolved. In other systems, MAPK activation leads to a nucleosomal response at IEG loci that involves phosphorylation and acetylation of serine and lysine residues respectively in the histone H3 NH_2_-terminal tail (Clayton et al., 2000; Dyson et al., 2005). Phosphorylated histone H3 in turn recruits scaffolding proteins such as 14-3-3 family members, downstream transcriptional regulators and chromatin remodelling factors to bring about gene expression changes. The absence of a consensus site for ERK1/2 phosphorylation in the NH_2-_terminus of histone H3 would suggest that its phosphorylation is not directly via ERK1/2 but a downstream kinase. A candidate for this activity is the family of nuclear-localised mitogen and stress activated kinases (MSK1/2), which have been reported to mediate histone H3 phosphorylation (Duncan et al., 2006; Soloaga et al., 2003). MSKs are nuclear-localised kinases comprising N- and C-terminal kinase domains separated by a flexible linker peptide. After an initial phosphorylation event by upstream MAPK including ERK1/2, MSKs undergo autophosphorylation, leading to full activation (Malakhova et al., 2009; McCoy et al., 2005). MSK1 and the highly homologous kinase MSK2 are both expressed in the heart and are activated in response to hypertrophic stimuli suggesting a potential role in hypertrophic remodelling (Deak et al., 1998a; Markou et al., 2004, 2009). A functional role of MSK in the heart and remains to be demonstrated however with studies thus far being compromised by the highly non selective inhibitors of MSK employed (Markou et al., 2009; Naqvi et al., 2012). Indeed, inhibitors used show equal efficacy on other kinases important in cardiomyocyte stress responses including RSK2, PKC isoforms and S6 kinase (Markou et al., 2009; Naqvi et al., 2012). Delineating whether MSK plays a role in cardiomyocyte responses and whether it contributes to the induction of the IEG response in these cells via a nucleosomal response remains key to understanding how cardiomyocytes respond to hypertrophic stimuli.

Here, we determined that MSK1/2 activated downstream of ERK1/2 was required for the cardiomyocyte hypertrophic response both in vitro and in vivo. MSKs elicits this function through promoting the phosphorylation of histone H3 S28 (pH3S28), which in turn recruits AP-1 factors, c-FOS and c-JUN along with the chromatin remodeller BRG-1 to IEG promoters, inducing transcription. Notably, the activation and role of the ERK1/2-MSK1/2-pH3S28 molecular axis was conserved in human samples demonstrating the relevance to human disease.

Together our data identify a key and missing component in the signalling pathway that transduces the activation of GPCRs by pathological pro-hypertrophic mediators in cardiomyocytes to the induction of hypertrophic remodelling.

## Results

### ERK1/2 activity is required for the induction of cardiomyocyte hypertrophic gene expression by ET-1 in vitro

MAPK signalling pathways, IEG induction and AP-1 transcription factor engagement contribute to the hypertrophic remodelling of cardiomyocytes (Archer et al., 2017). To examine the mechanisms underlying induction of IEG expression, the signalling pathways involved and their relationship to cellular hypertrophic responses were analysed in in vitro and in vivo models following acute exposure to established inducers of pathological hypertrophy.

Consistent with findings from our laboratory and elsewhere, application of the GPCR agonist endothelin-1 (ET-1) for 24 h stimulated a classical hypertrophic response in neonatal rat ventricular myocytes (NRVMs) that was associated with increases in mRNA levels of *Anf/Nppa* and Bnp/*Nppb*, in cell size and in the number of cells positive for perinuclear Anf protein (Figure 1A-C and S1A). Alongside this hypertrophic response, the expression of IEGs including *c-Fos* and *c-Jun*, was rapidly upregulated in ET-1 stimulated NRVMs (Archer et al., 2017) (Figure 1D, Figure S1B). These effects of ET-1 on both induction of hypertrophy and IEGs were prevented by inhibition of the MAPK pathway with PD184352 (PD, an inhibitor of the direct upstream kinase of ERK1/2, MEK1/2; Figure S1C) (Archer et al., 2017; Heineke and Molkentin, 2006) (Figure 1A-D). The efficacy of PD in preventing ERK1/2 activation (and hence MAPK pathway activation) was confirmed by immunoblotting, which showed a loss of the phosphorylated active form of ERK, pERK1/2, which was elevated in ET-1 stimulated NRVMs (Figure S1D). Baseline pERK1/2 in non-stimulated cells was also decreased following PD (Figure S1D).

**Figure 1:**
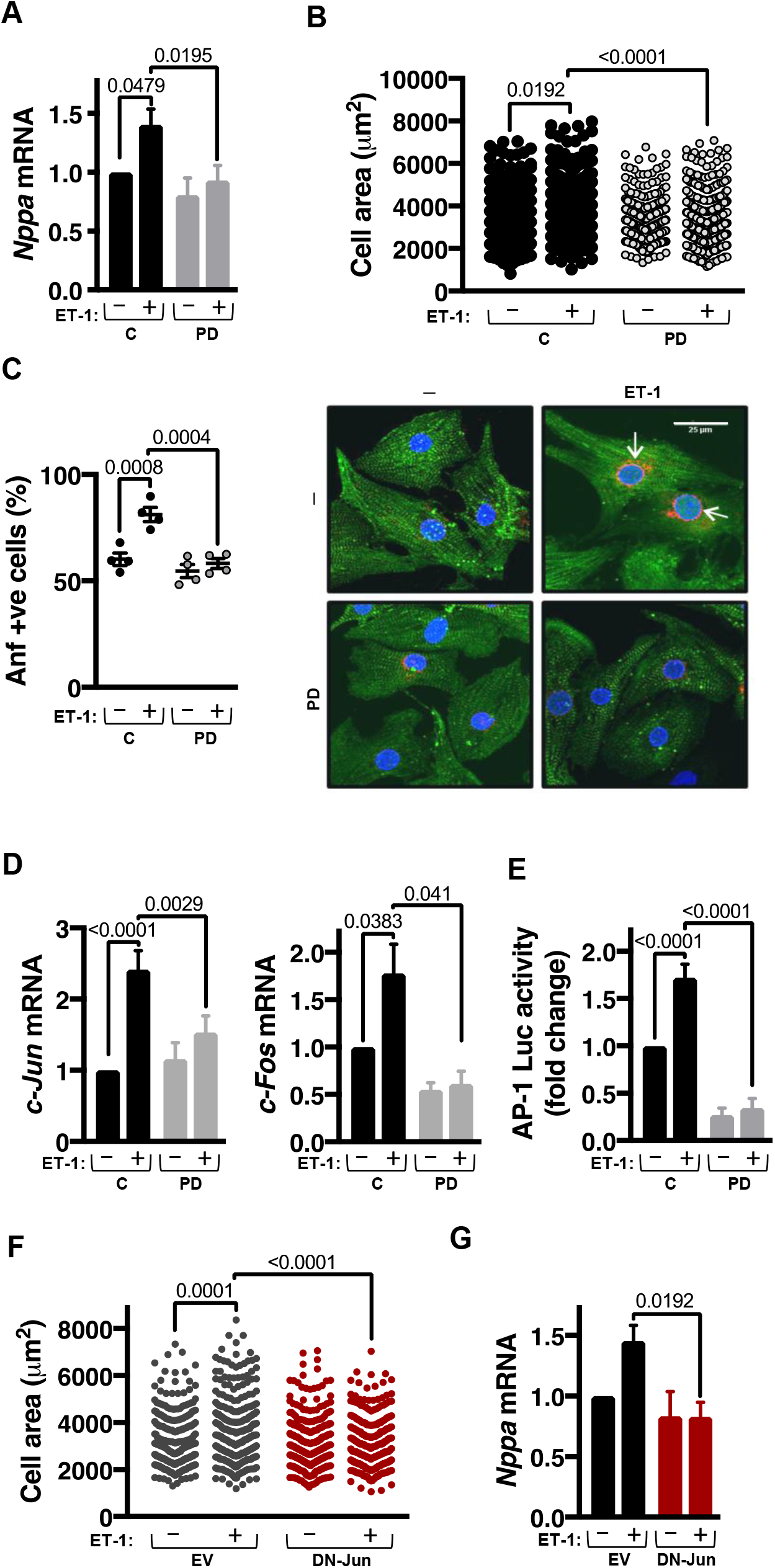
Pharmacological and molecular suppression of ERK signalling attenuates IEG induction and hypertrophy in vitro. **A**. RT-qPCR gene expression analysis for *Nppa/Anf* mRNA in NRVMs +/- 30 min pre-treatment with PD184352 (PD) +/- 15 min ET-1. N=5 biological replicates (defined as 5 different NRVM preparations from different litters), n=3 technical replicates (repeated reactions from the same sample). **B**. Cell area (µm^2^) as a measure of cardiomyocyte hypertrophy in NRVMs +/- PD +/- ET-1. N=4, pseudoreplicates (*<u>n</u>*) (defined as repeated measures of different cells or regions of interest from the same sample) =80-125. **C**. Immunostaining for Anf (red), sarcomeric alpha-actinin (α-Act) (green) and nuclear stained with DAPI (blue) in NRVMs +/- PD +/- ET-1. Quantification (left) represents individual mean data points for N=4, representative images (right). The scale bar represents 25 μm. **D**. RT-qPCR analysis of immediate early genes (IEGs) *c-Jun* and *c-Fos* mRNA expression in NRVMs +/- PD +/- ET- 1. For *c-Jun* (Left), N=4, n=3. For *c-Fos* (Right), N=3, n=3. **E**. AP-1 luciferase assay as a readout of AP-1 transcriptional activity in NRVMs +/- PD +/-ET-1 relative to untreated. For C and C ET-1, N=8, n=3. For PD and PD ET-1,N=3, n=3. **F**. Cell area (µm^2^) measurements for NRVMs infected with either control or DN-Jun (dominant negative, kinase dead) adenoviral vectors +/- ET-1. Individual data points are represented by N=4, *<u>n</u>*=80-100. **G**. RT-qPCR analysis of *Nppa*/Anf mRNA expression in NRVMs +/- DN-Jun +/- ET-1, N=4, n=3.

Consistent with the role of MAPK signalling to AP-1 in hypertrophy induction, a MAPK-dependent increase in activity of a luciferase-based AP-1 reporter was observed in ET-1 stimulated NRVMs (Figure 1E). Further, inhibition of AP-1 activity by adenoviral-mediated expression of dominant negative Jun (DN-Jun) abrogated the ET-1 stimulated increases in cell surface area and *Nppa* mRNA (Figure 1F-G).

Together, these data confirm that a pathway involving ERK1/2, IEG induction and AP-1 activity is engaged and required for the immediate early hypertrophic response to ET-1.

### Endothelin-1 stimulates ERK-dependent phosphorylation of Histone H3 Serine 28

The nucleosomal response – a term coined to link histone phosphorylation and IEG induction - is reported to be indissociable from ERK1/2 activity in a number of cell contexts. We therefore determined whether this nucleosomal response was engaged during ET-1 stimulated IEG induction in NRVM. Levels of histone H3 phosphorylated at Ser 10 and Ser 28 (pH3S10 and pH3S28) were quantified in histones acid-extracted from NRVMs exposed to ET-1. Consistent with the time-course of *c-Fos* induction, ET-1 promoted a significant increase in H3S28 phosphorylation at 10 min, which was not increased further at 30 min post ET-1 application (Figure 2A). Notably, the ET-1-stimulated increase in H3S28 phosphorylation was ERK pathway dependent, as shown by the loss of the response to ET-1 in cells treated with PD (Figure 2A). Neither ET-1 nor PD significantly affected H3S10 phosphorylation (Figure 2A).

**Figure 2:**
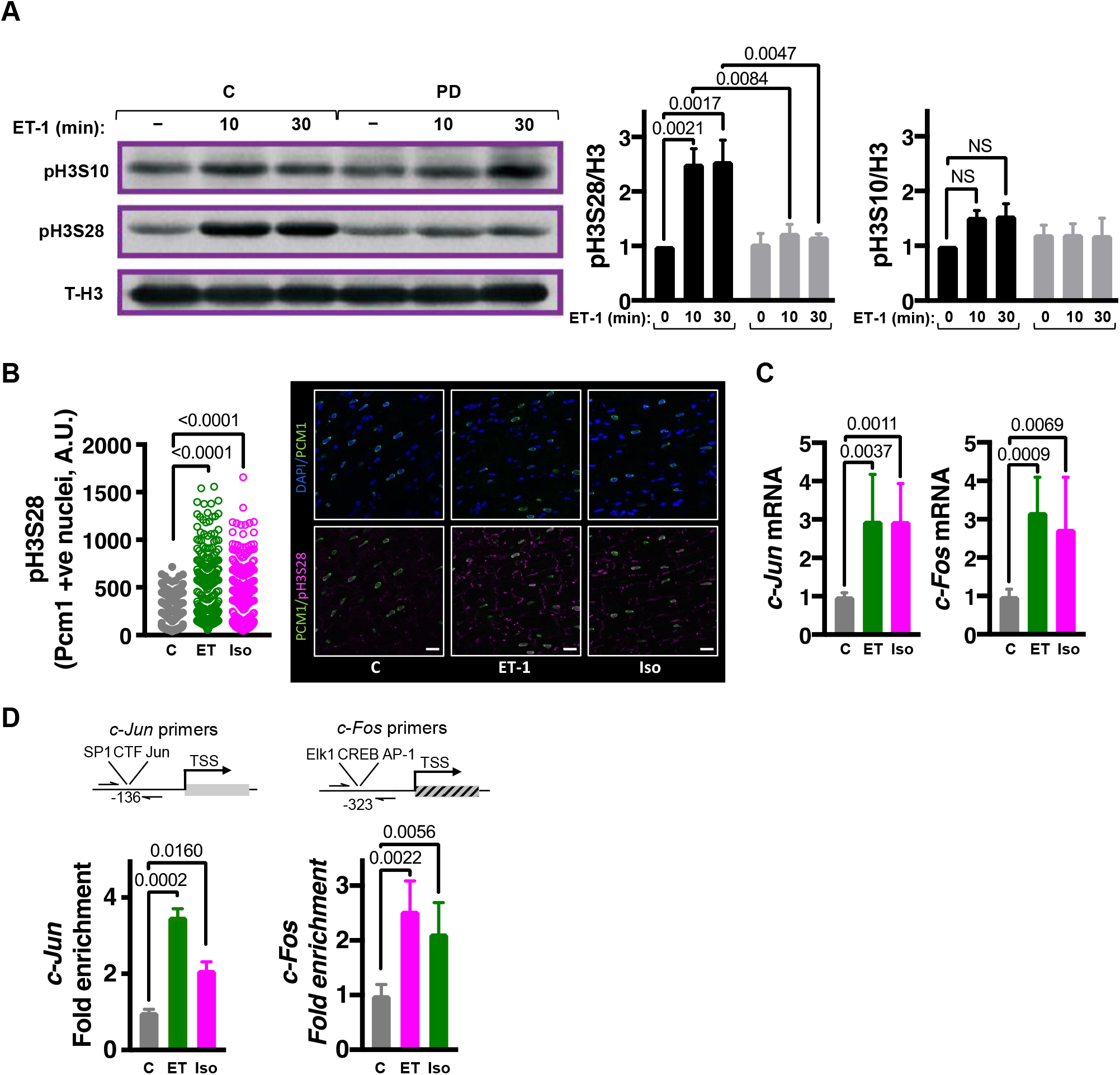
Neurohumoral signalling-induced ERK1/2 activation results in histone H3S28 phosphorylation at IEG promoters. **A**. Immunoblot analysis of pH3S10 and pH3S28. Left: Representative immunoblots from 1 NRVM preparation probing for pH3S10 and pH3S28 in acid extracted histones from NRVMs exposed to ET-1 for 0, 10 and 30 min in the presence or absence of PD. Right: Levels of phosphorylated histone normalised to total H3 are shown. For pH3S10, N=5. For pH328, N=3. **B**. Confocal immunofluorescence analysis of pH3S28 in cardiomyocytes in ventricular cardiac sections prepared from rats infused with ET-1 or Iso for 15 min. Cardiomyocyte nuclei were demarcated by pericentriolar material 1 (Pcm-1; in green) perinuclear staining. Nuclei are stained with DAPI (blue) and pH3S28 in magenta. Scale bar = 25 µm. The plot (left) shows quantification of nuclear pH3S28 in Pcm-1 positive nuclei. N=4, *<u>n</u>*=80-125. **C**. RT-qPCR gene expression analysis of IEGs *c-Jun* and *c-Fos* mRNA in left ventricular tissue from Wistar rats administered with ET-1 or Iso through jugular vein infusion and sacrificed 15 min later. C, N=7, ET-1, N=8, Iso, N=6, n=3. **D**. Chromatin immunoprecipitation-qPCR (ChIP-qPCR) analysis of pH3S28 at IEG promoters, *c-Jun* and *c-Fos* in adult male Wistar rats that were administered ET-1 or Iso through jugular vein infusion and sacrificed 15 min later. Top: schematic for the site of ChIP primer amplification relative to the transcription start sites. Below: quantification of enrichment compared with control (untreated) rats. N=3, n=3.

The stimulation of H3S28 phosphorylation by hypertrophic agonists was next measured in vivo. To this end, cardiomyocyte H3S28 phosphorylation was analysed in adult rats subsequent to 15 min infusion with sub-pressor levels of ET-1 via the jugular vein (Archer et al., 2017; Beyer et al., 1994; Dyson et al., 2005). Responses to the synthetic β-adrenergic agonist isoproterenol (Iso), which is often used chronically to induce pathological cardiac remodelling was also determined (Boluyt et al., 1995; Liu et al., 2009). Both ET-1 and Iso induced a significant increase in pH3S28 in cardiomyocyte nuclei (demarcated by Pcm-1 staining) in heart sections from rats infused with ET-1 and Iso (Figure 2B). Concurrent with increased pH3S28 in this in vivo model, mRNA expression of IEGs was induced following 15 min stimulation (Figure 2C and S2A). Supporting the pro-hypertrophic effect of these infusions with ET-1 or Iso, expression of the classic hypertrophy-associated foetal gene programme, including *Nppa/Anf* and *Nppb/Bnp* was also induced in these animals (Figure S2B).

To probe whether IEG promoters were phosphorylated at histone H3 during this intervention, chromatin immunoprecipitation (ChIP) experiments were performed. Chromatin was precipitated using antibodies against phosphorylated H3S28 and product detected by qPCR using primer pairs targeting the c-*Jun* and c-*Fos* promoters, as shown in the cartoons (Figure 2D). Notably, consistent with the increase in phosphorylated H3S28 detected by immunoblotting and immunofluorescence staining of bulk histones, ChIP enrichment for pH3S28-associated IEG promoters was substantially increased following 15 min ET-1 or Iso infusion (Figure 2D).

### MSK1/2 is activated following ET-1 stimulation in an ERK1/2 dependent manner and is required for IEG expression

Although ERK activation is required for ET-1 dependent phosphorylation of histone H3 at S28, the absence of consensus sequences for its phosphorylation of H3 would suggest that it is not the responsible histone-kinase. Rather, ERK activates a downstream effector kinase that in turn phosphorylates histone H3. A candidate for this action is the mitogen and stress-activated kinase (MSK1/2), which has been shown in other tissue contexts to phosphorylate H3S28 (Soloaga et al., 2003). While MSK1/2 activation following hypertrophic stimulation is reported in cardiomyocytes, it has not been shown to play a role in histone phosphorylation and IEG induction. To establish whether MSK1/2 was engaged during ET-1 stimulation of cardiomyocytes and whether ERK activity was involved, levels of MSK1 phosphorylated at S376, a phosphorylation event required for activity, were analysed by immunoblotting of lysates prepared from NRVM stimulated with ET-1-stimulated ± PD (Figure 3A). As shown in the immunoblot and accompanying densitometric analysis, ET-1 stimulated an increase in pMSK that was sensitive to MEK/ERK pathway inhibition with PD (Figure 3A). Importantly, using the in vivo model described earlier, an increase in pMSK was also detected in cardiomyocyte nuclei in heart sections from rats infused with ET-1 or Iso for 15 min (Figure 3B). Together, these experiments indicated that hypertrophic agonists stimulate a rapid phosphorylation of MSK in cardiomyocytes that is dependent on activation of MEK/ERK signalling.

**Figure 3:**
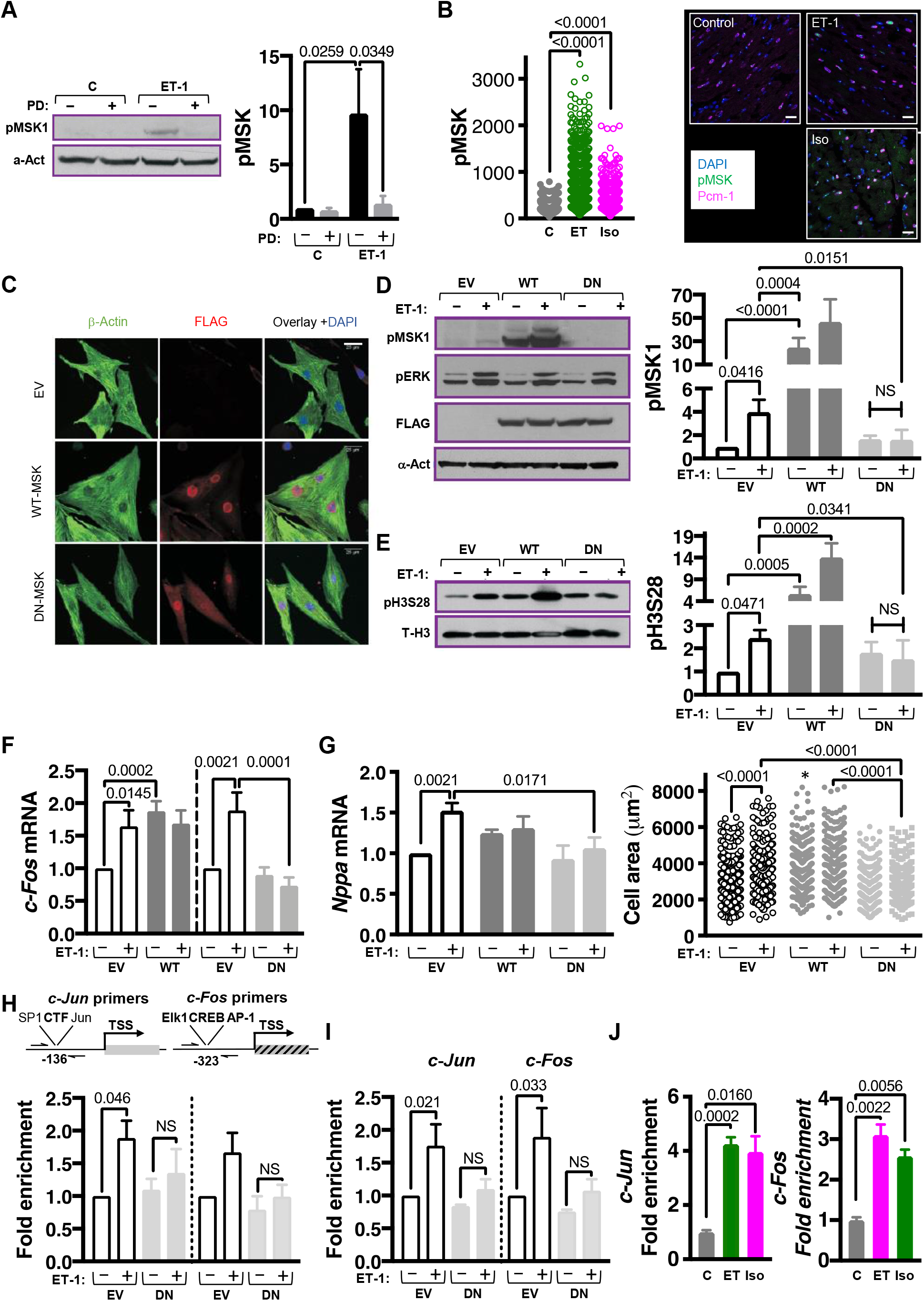
Activated MSK is required for histone H3S28 phosphorylation, recruitment of Brg1 to chromatin and IEG induction in cardiomyocytes. **A**. Immunoblot showing levels of phosphorylated (activated) MSK in NRVMs +/- PD +/- ET-1, normalised to αAct as a loading control. Left: Representative immunoblot. The α-Actinin blot is the same as shown in Fig S1D and pMSK was probed on the same blot. Right: Quantification of pMSK relative to control vehicle treated cells. N=5. **B**. Confocal immunofluorescence analysis of pMSK in cardiomyocytes in ventricular cardiac sections prepared from rats infused with ET-1 or Iso for 15 min. Cardiomyocyte nuclei were demarcated by pericentriolar material 1 (Pcm-1; in magenta) perinuclear staining. Nuclei are stained with DAPI (blue) and pMSK in green. Left: Quantification of nuclear pMSK in Pcm-1 positive nuclei. N=4, *<u>n</u>*=200-400. Right: Confocal images of heart sections from animals treated as indicated. Scale bar = 20 µm. **C**. Representative confocal images of immunostained NRVMs showing expression of FLAG-tagged WT-MSK and DN-MSK adenoviruses (AdV). Nuclei are stained with DAPI (blue), Beta-Actin in green and FLAG-tagged MSK in red. **D**. Immunoblotting for pMSK, pERK and FLAG-tagged MSK AdV in NRVMs infected with either empty vector (EV), WT-MSK1 AdV or DN-MSK1 AdV and treated +/-15 min with ET-1, normalised to α-Act as a loading control. Left: Representative immunoblot. Right: Quantification of immunoblot, relative to EV. N=5. **E**. Immunoblotting for phosphorylated histone H3S28 in NRVMs infected with either empty vector (EV), WT-MSK1 AdV or DN-MSK1 AdV treated +/- 15 min with ET-1, normalised to total Histone 3 (T-H3) as a loading control. Left: Representative immunoblot. Right: Quantification of immunoblot data. N=6. **F**. Effect of DN-MSK expression on *c-Fos* expression in NRVMs treated with ET-1 for 10 min. *c-Fos* expression was determined by RT-qPCR. Data is presented relative to empty vector. For WT-MSK data (left-hand side), EV ctrl and WT-MSK ctrl, N =10, EV ET-1 and WT-MSK ET-1, N=6, n=3. For DN-MSK data (right-hand side), N=6, n=3. **G**. Analysis of hypertrophic responses in NRVMs infected with EV or DN-MSK1 AdV treated +/- ET-1 for 24 h. Left: RT-qPCR expression analysis of *Nppa*/*Anf* mRNA in NRVMs. Data is presented relative to EV untreated cells. For EV ctrl, EV ET-1, WT-MSK ctrl and WT-MSK ET-1, N=8, n=3. For DN-MSK ctrl and DN-MSK ET-1, N=6, n=3. Right: Cell area (μm^2^) as a measure of hypertrophy in NRVMs. N=4, *<u>n</u>*=50-80. **H**. ChIP-qPCR analysis for pH3S28 abundance at *c-Jun* (left) and *c-Fos* gene promoter regions in NRVMs infected with EV or DN-MSK1 AdV +/- ET-1 for 10 min. Top: schematic for the site of ChIP primer amplification relative to the transcription start sites. Below: quantification of enrichment compared with EV AdV untreated NRVMs. For *c-Jun* ChIP data (left-hand side), N=4, n=3. For *c-Fos* ChIP data (right-hand side), N=3, n=3. **I**. ChIP-qPCR analysis for Brg1 enrichment at *c-Jun* (left) and *c-Fos* gene promoter regions in NRVMs infected with EV or DN-MSK1 AdV +/- ET-1 for 10 min. Quantification of Brg1 enrichment at the *c-Jun* (Left) and *c-Fos* (Right) promoters compared with EV AdV untreated NRVMs. For *c-Jun* ChIP data (left-hand side), N=4, n=3. For *c-Fos* ChIP data (right-hand side), N=3, n=3. **J**. ChIP-qPCR for Brg1 at the *c-Jun* and *c-Fos* gene promoters (left-right) in left ventricular tissue from adult male Wistar rats that were administered ET-1 or Iso through jugular vein administration and sacrificed 15 min later. N=3, n=3.

Experiments were next conducted to investigate the functional role of MSK in the nucleosomal response to ET-1 in cardiomyocytes. Owing to the lack of availability of specific and sensitive pharmacological agents targeting MSK1/2 (Bain et al., 2007), a molecular approach was adopted to manipulate its activity. To this end, adenoviruses were first employed. MSK1 signalling was enhanced through overexpression of wild-type MSK1 (WT-MSK), whilst endogenous MSK1 was inhibited by expression of a D565A kinase dead mutant of MSK1 that acts in a dominant–negative fashion (DN-MSK1) (Deak et al., 1998a). Owing to shared regulatory mechanisms of MSK1 and MSK2, DN-MSK1 would be expected to inhibit activity of both kinases, overcoming possible redundancy (Zhong et al., 2001). Immunofluorescence analysis demonstrated that adenovirally-expressed FLAG-tagged WT- and DN-MSK1 were both localised to the nucleus in NRVMs (Figure 3C). Overexpression of the WT and DN-MSK1 proteins at equivalent levels was confirmed by immunoblotting using an antibody directed against their NH_2_-terminal FLAG epitope (Figure 3D). Overexpression of WT-MSK1 produced an increase in baseline MSK1 phosphorylation whilst DN-MSK1 prevented the activation of endogenous MSK1 in response to ET-1, consistent with a dominant negative effect (Figure 3D). Neither overexpression of WT-MSK1 or DN-MSK1 affected ERK1/2 activation by ET-1, indicating that effects of these strategies to modify MSK activity are not mediated via altered ERK1/2 activity but by MSK1 itself (Figure 3D). The consequences of WT and DN-MSK1 expression on ET-1 stimulated histone H3S28 phosphorylation were next measured. Notably, both ET-1 and overexpression of WT-MSK1 elevated levels of pH3S28 in NRVMs, whereas expression of DN-MSK1 prevented the ET-1 stimulated increase in pH3S28 (Figure 3E), thereby demonstrating the requirement for MSK activity for H3S28 phosphorylation during the hypertrophic response to ET-1.

Whether the effect of WT and DN-MSK on MSK activation in ET-1 stimulated NRVM translated to an effect on IEG induction and hypertrophic responses was next examined (Figure 3F-G). NRVM expressing WT-MSK expression exhibited a significant elevation in *c-Fos* expression compared to control, which was not increased further by 10 min exposure to ET-1 (Figure 3F). Consistent with its effects on kinase activation and H3S28 phosphorylation, DN-MSK1 expression significantly inhibited ET-1 stimulated *c-Fos* induction. WT-MSK expression also had a significant effect on cell hypertrophy, stimulating an increase in cell area in the absence of ET-1 (Figure 3G). WT-MSK1 did not promote a significant increase in *Nppa* mRNA. Notably DN-MSK1 expression suppressed hypertrophic responses in NRVM exposed to ET-1 for 24 h, with a significant inhibition of the ET-1 stimulated increases in *Nppa* mRNA and in cell size observed when compared to controls (Figure 3G). Further supporting the role of MSK in histone H3 S28 phosphorylation, IEG induction and hypertrophy, abrogation of ET-1 stimulated increases in H3S28 phosphorylation, *c-Fos* and *Nppa* mRNA expression were also observed in ARVM (Figure S3A-C).

Having validated that suppression of MSK activity via DN-MSK1 expression could prevent ET-1 stimulated phosphorylation of histone H3S28, *c-Fos* induction and hypertrophic responses, we next probed the requirement for MSK for ET-1 stimulated phosphorylation of H3S28 at IEG promoters. To this end, ChIP experiments were carried out in NRVMs expressing DN-MSK1 or empty vector control that were exposed to ET-1 or vehicle for 10 min. Importantly, DN-MSK1 prevented the ET-1-stimulated increase in pH3S28 specifically at IEG promoters (Figure 3H). In these experiments, the ET-1 stimulated ChIP enrichment of c-*Jun* and c-*Fos* promoters was lost in NRVM overexpressing DN-MSK1 (Figure 3H).

To complement the experiments using DN-MSK, the requirement of MSK activity for ET-1 induction of *c-Fos, Msk1* mRNA expression was tested by knocking down its expression using small interfering RNA (siRNA). Using this approach, ∼60 % reduction in *Msk1* mRNA was achieved compared to NRVM transfected with scrambled control siRNA (Figure S3C). In Msk1 siRNA knockdown (siMsk1) NRVMs, ET-1 stimulated induction of *c-Fos* was significantly blunted after 10 min exposure (Figure S3D). Moreover, siMsk1 prevented ET-1 stimulated induction of hypertrophic gene expression (Figure S3E).

Together, these data generated using siRNAs and adenoviruses to manipulate both MSK activity and expression not only indicate that H3S28 is a substrate for MSK1 but also that MSK1 is the kinase responsible for the phosphorylation of H3S28 and induction of *c-Fos* expression in cardiomyocytes stimulated with ET-1. Moreover, our data shows that these events are required for the hypertrophic response of cardiomyocytes.

### MSK1-mediated phosphorylation of H3S28 recruits BRG1, a component of the SWI/SNF family of chromatin remodellers to IEG loci

Previous studies have reported a requirement for the BRG1 (encoded by gene *SMARCA4*) component of the BAF (BRG1/BRM-associated factor 60; BAF60) ATP-dependent SWI/SNF chromatin remodelling complex in induction of IEG expression and Myh isoform switching in pathological cardiac remodelling (Deak et al., 1998b; Hang et al., 2010). The role of Brg1 in IEG induction and hypertrophic remodelling was therefore tested. Consistent with previous studies, knock down of Brg1 with siRNA prevented isoform switching between *Myh6* and *Myh7*, associated with induction of pathological hypertrophy, and abrogated *c-Fos* induction in ET-1 stimulated NRVMs (Figure S3F-G). Having shown the involvement of Brg1 in ET-1 responses in NRVM, whether phosphorylation of H3S28 affected Brg1 occupancy at IEG promoters was next investigated. To this end, ChIP experiments were performed on NRVMs expressing DN-MSK1 ± ET-1, as in Figure 3H but using an antibody directed Brg1. Notably, as shown by greater ChIP enrichment, ET-1 stimulated increases in association of Brg1 with both the *c-Jun* and *c-Fos* promoters in NRVMs (Figure 3I). Brg1 recruitment was however prevented in DN-MSK expressing NRVMs. Further supporting these in vitro data, Brg1 was also enriched at the *c-Jun* and *c-Fos* promoters in chromatin prepared from hearts from Wistar rats infused with ET-1 or Iso for 15 min (Figure 3J).

Collectively, these data demonstrate that MSK-mediated phosphorylation of histone H3S28 at the promoter regions of IEGs is a necessary event for the recruitment of the chromatin remodelling complex to these promoters required for IEG expression and hypertrophic remodelling subsequent to GPCR stimulation.

### MSK1/2 expression is required for the hypertrophic response in vivo

The data above indicate a role for MSK in the induction of hypertrophic responses in vitro and that the MSK/pH3S28/Brg1/IEG pathway is acutely engaged following neurohumoral stimulation in vivo. The requirement for MSK1/2 for activation of this pathway and induction of hypertrophy was next examined in vivo using a mouse model in which both alleles of *Msk* had been deleted by homologous recombination (*Msk1/2*^-/-/-/-^; *Msk* KO) (Wiggin et al., 2002a). Specifically, wild type (WT) and *Msk* knockout (KO) mice were subjected to osmotic mini-pump infusion of Iso for one week, after which the activation of the MSK pathway and hypertrophy induction was measured. During this one week infusion, the initial adaptive remodelling associated with pathological hypertrophy would be expected.

The activation of the MSK/pH3S28/Brg1/IEG pathway axis during the one week Iso infusion and how it was affected by loss of Msk1/2 was first assessed. As expected for the *Msk* KO mice, *Msk1* and *Msk2* transcripts were absent in heart tissue in this mouse (Figure 4A). *Msk1* and *Msk2 mRNA levels* were also measured in the iso infused WT mice and found to be significantly upregulated after 1 week (Figure 4A). Notably, Iso-infused WT mice exhibited significant increases in expression of IEGs including *c-Fos* and *c-Jun* and in line with its role in IEG induction of *Smarca4* (the gene encoding Brg1) (Figure 4B-C). Consistent with the importance of MSK1/2 in the IEG response, Iso induction of *c-Fos, c-Jun* and *Smarca4* expression was reduced in these mice when compared to similarly treated WT controls (Figure 4B-C). Levels of nuclear pMSK and pH3S28 (demarcated by Pcm-1 or Nesprin perinuclear staining (Thienpont et al., 2017) in cardiomyocytes in sections prepared from Iso-infused WT and KO animals were also significantly lower in KO animals than WT at baseline and were not affected by Iso, indicating inactivation of the MSK/pH3S28 pathway (Figure 4D-E). While significantly higher than in KO at baseline, no significant effect of Iso on pMSK or pH3S28 was however observed in WT cardiomyocytes.

**Figure 4:**
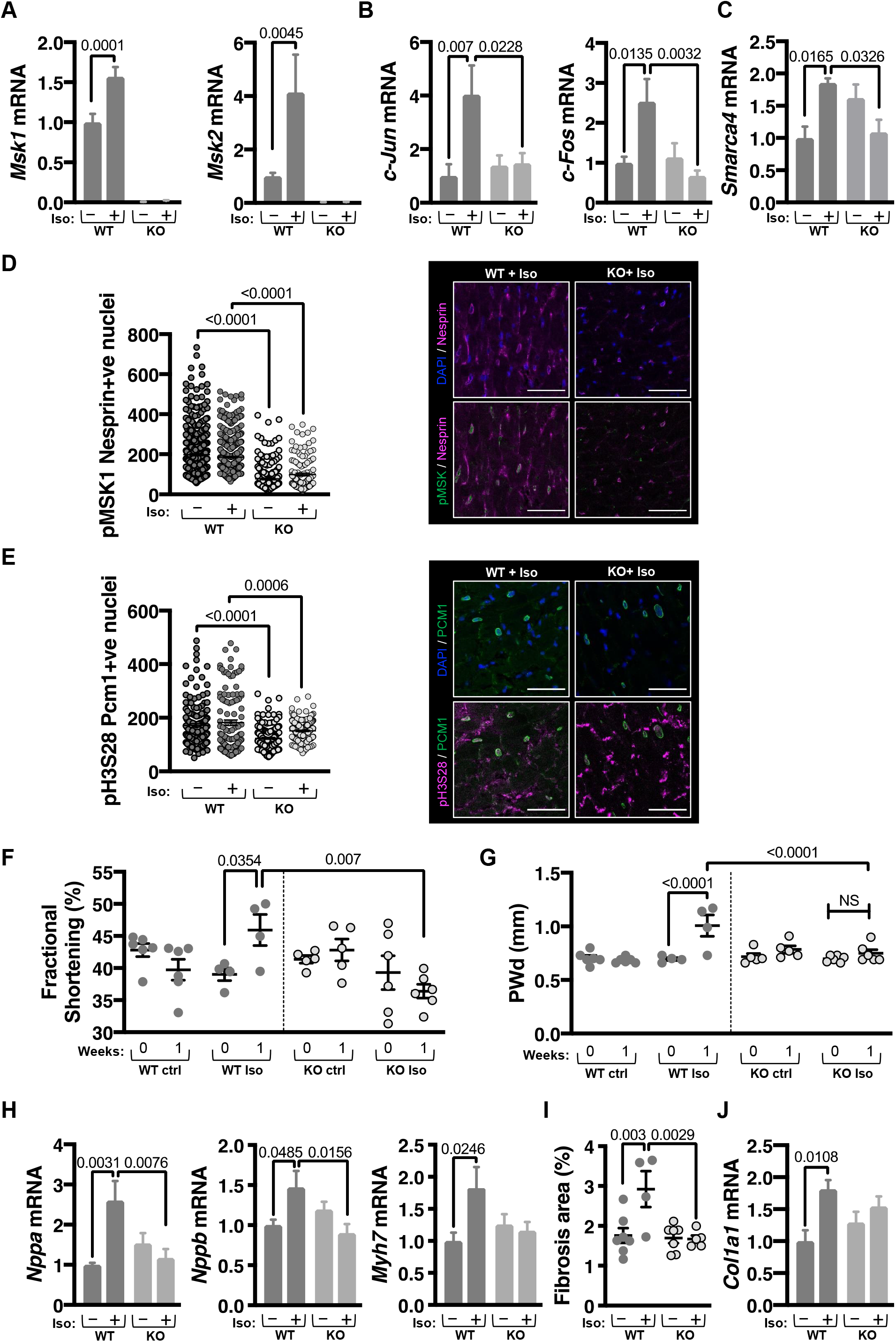
Genetic MSK inhibition attenuates IEG activation and cardiomyocyte hypertrophy in vivo. **A**. RT-qPCR analysis of *Msk1* (Left) and *Msk2* (Right) mRNA expression in left ventricle from *Msk1/2* KO mice and wild type littermates +/- Iso infusion for 1 week. WT ctrl, N=7, WT Iso, N=4, KO ctrl, N=5, KO Iso, N=8, n=3. **B**. RT-qPCR analysis of expression of IEGs *c-Jun* (Left) and *c-Fos* (Right) in left ventricle from *Msk1/2* KO mice and wild type littermates +/- Iso infusion for 1 week. WT ctrl, N=5, WT Iso, N=4, KO ctrl, N=5, KO Iso, N=5, n=3. **C**. RT-qPCR analysis of *Smarca4/Brg1* mRNA expression in left ventricle from *Msk1/2* KO mice and wild type littermates +/- Iso infusion for 1 week. WT ctrl, N=5, WT Iso, N=4, KO ctrl, N=5, KO Iso, N=5, n=3. **D**. Immunostaining for pMSK in cardiomyocyte nuclei in left ventricular cardiac sections in *Msk1/2* KO mice and wild type littermates +/- Iso infusion for 1 week. Cardiomyocyte nuclei are demarcated with Nesprin. Left: Quantification of pMSK in Nesprin+ve nuclei. Right: Representative immunostaining images for pMSK (green), Nesprin (red) and nuclei are stained with DAPI (blue). N=4, *<u>n</u>*=30-160. Scale bar = 50 µm. **E**. Immunostaining for pH3S28 in cardiomyocyte nuclei in left ventricular cardiac sections in *Msk1/2* KO mice and wild type littermates +/- Iso infusion for 1 week. Cardiomyocyte nuclei are demarcated with Pcm-1. Left: Quantification of pH3S28 in Pcm-1+ve nuclei. Right: Representative immunostaining images for pH3S28 (red), Pcm-1 (green) and nuclei are stained with DAPI (blue). N=4, *<u>n</u>*=30-160. Scale bar = 50 µm. **F**. Fractional shortening in *Msk1/2* KO mice and wild type littermates at baseline (Iso=0) and +/- Iso infusion for 1 week (Iso=1), derived from 2D echocardiography data. WT ctrl, N=6, WT Iso, N=4, KO ctrl, N=5, KO Iso, N=6. **G**. Posterior wall dimension in diastole in *Msk1/2* KO mice and wild type littermates at Iso=0 and +/- Iso=1, derived from 2D echocardiography data. WT ctrl, N=6, WT Iso, N=4, KO ctrl, N=5, KO Iso, N=6. **H**. RT-qPCR analysis of the markers of pathological hypertrophy *Nppa*/Anf, *Nppb*/Bnp and *Myh*7 mRNA expression in left ventricle from *Msk1/2* KO mice and wild type littermates +/- Iso infusion for 1 week. N=5, WT Iso, N=4, KO ctrl, N=5, KO Iso, N=5, n=3. **I**. Quantification of left ventricular interstitial fibrosis, measured as percentage (%) area of extracellular matrix from Picro Sirius Red staining in left ventricular tissue from *Msk1/2* KO mice and wild type littermates +/- Iso infusion for 1 week. WT ctrl, N=7, WT Iso, N=4, KO ctrl, N=7, KO Iso, N=5, n=3. **J**. RT-qPCR analysis of *Col1a1* mRNA expression in left ventricular tissue from *Msk1/2* KO mice and wild type littermates +/- Iso infusion for 1 week. WT ctrl, N=5, WT Iso, N=4, KO ctrl, N=5, KO Iso, N=5, n=3.

We tested whether upregulation of IEGs, *Smarca4* and *Msk*1/2 was a feature of other models of hypertrophic remodelling. In line with these findings in Iso infused mice, an upregulation of IEGs, *Smarca4* and of *Msk1/2* was detected in cardiomyocytes from rats subjected to constriction of the ascending aorta (AB) for 6 weeks (a model of pathological hypertrophy) (Thienpont et al., 2017) (Figure S4A). Indicative of a specific role of this IEG response to pathological cardiac remodelling, no such upregulation was detected in cardiomyocytes from rats subjected to treadmill training for 6 weeks, which exhibited physiological cardiac remodelling, with a similar degree of hypertrophy as the AB rats (Thienpont et al., 2017) (Figure S4A). *Msk1* and *2* mRNA levels were also increased in NRVMs after 24 h of ET-1 stimulation (Figure S4B).

Having confirmed the engagement of the MSK/pH3S28/Brg1/IEG pathway in Iso-infused mice, and the requirement for MSK1/2 for phosphorylation of H3S28 and induction of IEG, hypertrophic remodelling was assessed in these animals. Cardiac function and geometry were measured in vivo by 2D echocardiography prior to the start of the experiment (baseline) and after one week of Iso infusion. In line with the reported lack of an overt phenotype in *Msk* double KO mice, no differences in cardiac function, indicated by fractional shortening (FS) or of posterior wall thickness (PWd) were detected between control WT and KO animals at baseline (Figure 4F and S4C). Following one week Iso infusion, whereas control mice exhibited significant increases in PWd and FS, typical of the initial stages of pathological hypertrophic remodelling (Selvetella et al., 2004), these responses to Iso were absent in *Msk* KO mice (Figure 4F-G and S4C).

The hypertrophic response to Iso infusion was also assessed by RT-qPCR analysis of components of the foetal gene program. As would be expected, Iso infusion resulted in increased expression of *Nppa, Nppb* and *Myh7* in WT mice (Figure 4H). The expression of these hypertrophic markers was not however induced following Iso infusion in *Msk* KO mice, thereby supporting the requirement for Msk for the hypertrophic response observed in vitro.

Fibrosis is a feature of pathological hypertrophy, including following Iso infusion that contributes to disease progression. Histological analysis of LV tissue sections revealed a significant increase in interstitial fibrosis following Iso infusion in control mice that was absent in the *Msk* KO mice (Figure 4I and S4D). Supporting the histology analysis, *Col1a1* mRNA was substantially increased in Iso infused WT mice that was absent in similarly treated *Msk* KO mice (Figure 4J).

Fibrosis is a natural response to increased cardiomyocyte death in the myocardium. Given the association between cardiomyocyte viability and the activity of MAPK pathways, we assessed whether reduced cell death was contributing to the protective effects of Msk KO. TUNEL staining of heart sections revealed a significant Iso-dependent increase in cell death in WT mouse hearts that was not observed in *Msk* KO mice (Figure S4E). Baseline cell death was similar between WT and *Msk* KO mouse hearts. The expression of key anti- and pro-apoptotic mediators was analysed (Figure S4F and G). Expression of executioner caspases 3 and 9 (*Casp3* and *Casp9*) of apoptosis and of *Bax*, a pro-apoptotic BH3 only family member, were increased following Iso in WT mice, whereas their expression was not induced in the *Msk* KO mouse (Figure S4F). Notably, Bcl2, which exhibits anti-apoptotic activity, was increased in the *Msk* KO mouse following Iso infusion (Figure S4G).

Together these data show that consistent with that observed in vitro, MSK activity is required for the induction of hypertrophy in vivo, and that in its absence, hearts are protected from pathological insult.

### The MSK1/2/pH3S28/BRG1/IEG axis is engaged in human hypertrophic remodelling

Having shown an important role of MSK1/2 in IEG and hypertrophy induction in cellular models and in vivo in rodents, we analysed whether the MSK1/2/pH3S28/IEG axis was conserved in GPCR responses and hypertrophic remodelling in human. To this end, the activation of MSK and phosphorylation of H3S28 following application ET-1 and Iso was first analysed in acutely isolated ventricular cardiomyocytes from explanted non-failing donor hearts. The involvement of MAPK activity was also tested. Stimulation of human cardiomyocytes resulted in a significant increase in levels of pMSK and pH3S28 (Figure 5A-B). Consistent with our findings in rat ventricular cardiomyocytes, these phosphorylation events were abrogated by inhibition of the ERK1/2 pathway with PD.

**Figure 5:**
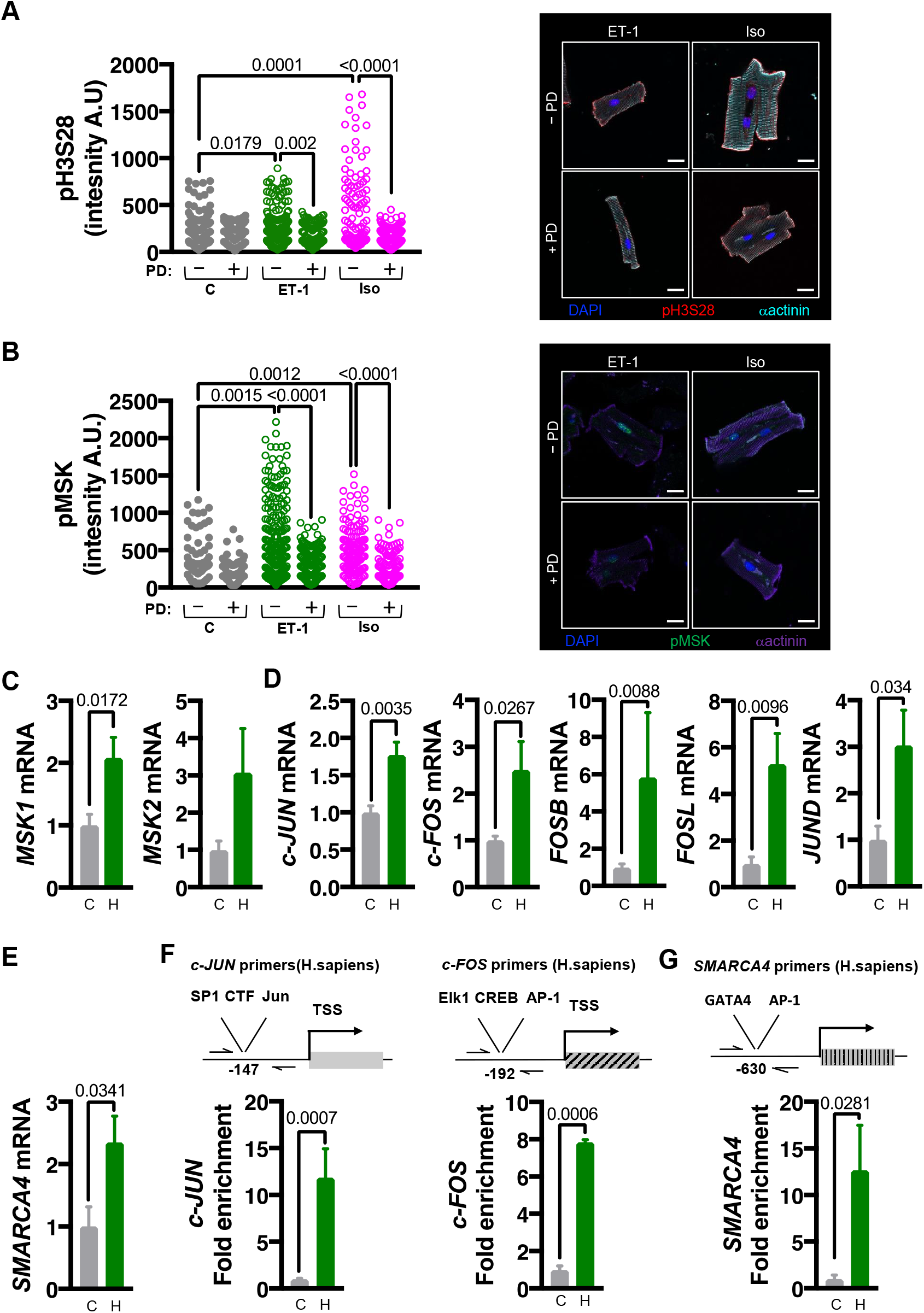
The MAPK-MSK-pH3S28 axis is conserved in the hypertrophic response in humans. **A**. Confocal immunofluorescence analysis of pH3S28 in isolated human donor cardiomyocytes treated for 15 min with ET-1 or Iso +/- PD. Left: Quantification of immunostaining of nuclear pH3S28. Right: Representative images of isolated cardiomyocytes stained for pH3S28 (red), α-Act (cyan), nuclei stained with DAPI (blue). N=4, *<u>n</u>*=12 - 123. Scale bar = 20 µm. **B**. Confocal immunofluorescence analysis of pMSK in isolated Immunostaining in isolated human donor cardiomyocytes treated for 15 min with ET-1 or Iso +/- PD. Left: Quantification of immunostaining of nuclear pMSK. Right: Representative images of isolated cardiomyocytes stained for pMSK (green), α-Act (purple), nuclei stained with DAPI (blue). N=4, *<u>n</u>*=13 - 136. Scale bar = 20 µm. **C-E**. RT-qPCR analysis of mRNA expression of indicated genes in human hypertrophic left ventricular tissue (H) compared with non-failing (C). In Figure 5C and 5E, C, N=5, H, N=4, n=3. In Figure 5D, C, N=4, H, N=4, n=3. **C**. RT-qPCR analysis of *MSK1* and *MSK2* mRNA expression. **D**. RT-qPCR analysis of expression of immediate early gene components of the AP-1 transcription factor. **E**. RT-qPCR analysis of *SMARCA4* (*BRG-1)*. **F**. ChIP-qPCR analysis for pH3S28 enrichment at the *c-JUN* and *c-FOS* promoters in Pcm-1 +ve cardiomyocyte nuclei from human hypertrophic left ventricular tissue (H) compared with non-failing (C). N=3, n=3. **G**. ChIP-qPCR for pH3S28 enrichment at the *SMARCA4/BRG-1* promoter in Pcm-1 +ve cardiomyocyte nuclei from human hypertrophic left ventricular tissue (H) compared with non-failing (C). N=3, n=3.

To gain insight into the involvement of MSK and IEG in human disease, the expression of MSK1/2, IEGs and early response gene target *SMARCA4* was compared between cardiomyocyte nuclei purified from healthy and hypertrophic human hearts. As in rodents, expression of MSK1/2, IEG components of the AP-1 transcription factor and SMARCA4 were substantially upregulated in hypertrophic cardiomyocytes (Figure 5C-E). We next analysed pH3S28 in the promoters of *c-FOS, C-JUN* and *SMARCA4* by ChIP. Notably, pH3S28 enrichment was observed at all three promoters in hypertrophic compared with healthy control hearts (Figure 5F-G).

Together, these data show conservation of the MSK pathway to IEG induction in human hearts and support our hypothesis that the MSK/pH3S28/IEG pathway is necessary to bring about the initial stages of the pathological hypertrophic response in cardiomyocytes (Figure 6).

**Figure 6:**
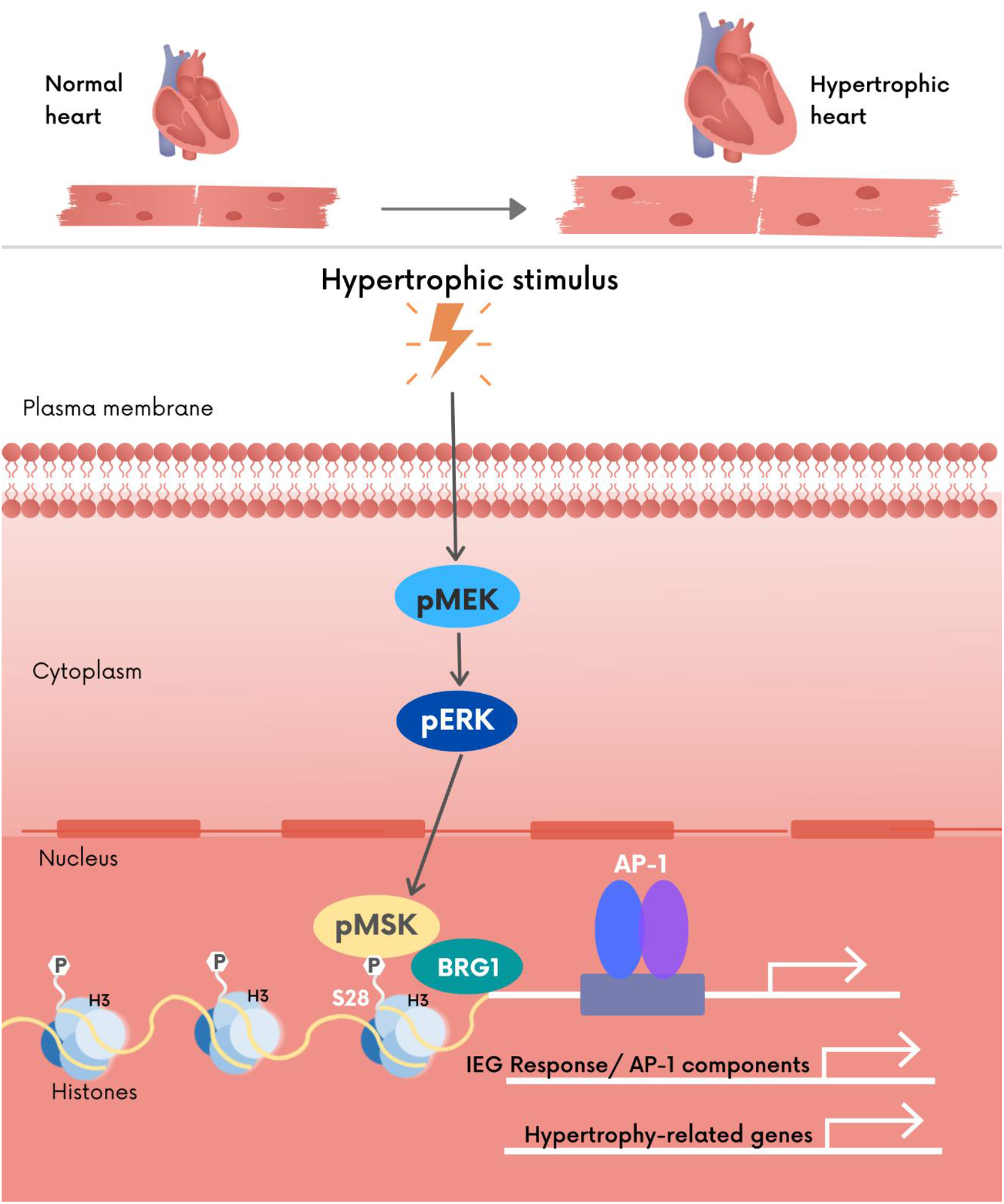
Summary cartoon of main findings of this study indicating pathway by which MSK couples GPCR activation with IEG induction during the cardiac hypertrophic response

## Discussion

MAPK regulation of immediate early gene activity and expression is key to stress-mediated induction of cardiac hypertrophic responses. Here we identified MSK1/2, a kinase activated downstream of ERK1/2, as being necessary for the initiation of IEG expression in response to pathological hypertrophic cues. MSK1/2 elicited this response through phosphorylation of histone 3 at Ser 28 allowing recruitment of the ATP-dependent chromatin remodeller, Brg1. In the absence of this response, hypertrophic gene expression and cardiac remodelling was attenuated. Notably, live-cell functional assays and analysis of post-mortem human hypertrophic hearts revealed that this pathway was conserved in humans. These data are summarised in the cartoon in Figure 6.

The hypertrophic response of the pathologically stressed myocardium is mediated through an extensive remodelling of the cardiomyocyte transcriptome (Selvetella et al., 2004; Song et al., 2012). Contributing to the initiation of this process as well as being required for its manifestation are AP-1 transcription factors, that comprise heterodimers of, but not restricted to, the proto-typical IEGs c-FOS and c-JUN (Chiu et al., 1988; Hess et al., 2004). Through their signal responsive phosphorylation, AP-1 are acutely activated in response to hypertrophic stimuli and upon phosphorylation associate with cognate response elements in the promoters of their encoding genes as well as numerous other targets, thereby promoting a rapid induction of their expression. Consistent with previous work from our laboratory and others and this established paradigm for AP-1 activity in the heart, we show here that the expression of these IEGs is increased within minutes of exposure of cells to hypertrophic agonists, both in vitro, and importantly in vivo (Archer et al., 2017). As revealed by use of dominant negative AP-1 components shown here and in previous studies, the induction of these early response genes, which include many transcription factors, is important in mediating later phases of the hypertrophic response (Hilfiker-Kleiner et al., 2006; Petrich et al., 2003).

### ERK1/2 in the heart

Activation of the ERK1/2 MAPK pathways is a conserved feature of many hypertrophic stressors ((Bueno et al., 2000)(20959622). This signalling cascade is activated following receptor engagement at the plasma membrane and culminates in induction of hypertrophic gene expression. As we show here and described elsewhere, expression of immediate early genes occurs in vitro and in vivo within minutes of exposure to hypertrophic agonist, and is the first transcriptional readout of MAPK activation. Indeed, we show that ERK1/2 activation mirrors the temporal profile of IEG induction. Moreover, in the absence of ERK activity, hormone stimulated AP-1 activation, IEG induction and expression of markers of hypertrophy in myocyte cultures is prevented in vitro and in vivo (Liu et al., 2016). Notably, while ERK and IEG activity peak proximal to the initiating stimulus, despite a lack of detectable activity subsequent to this peak, ERK inhibition prevents hypertrophic gene expression (Archer et al., 2017).

While we, as others, show an involvement of ERK1/2 and activation of IEGs in the mechanism underlying pathological hypertrophic remodelling, the role of ERK1/2 in hypertrophy in vivo is complex. Whereas in vivo overexpression of ERKs is not sufficient to induce a hypertrophic response, overexpression of its direct upstream kinase MEK induces a concentric hypertrophic response (Mutlak and Kehat, 2015). However, overexpression of the small GTPase protein Ras, which lies between GPCR activation and ERK activation, results in cardiomyopathy (Wu Guangyu et al., 2001). Recent elegant strategies involving conditional or tissue specific manipulation of each isoform to overcome embryonic lethality of the genetic KO now provide a clearer view of the contribution of this pathway to the cardiac hypertrophic response (Kehat et al., 2011; Purcell et al., 2007; Ulm et al., 2014). Loss of ERK2, which represents 50-70 % of ERK activity in the heart attenuates the initial compensatory phase of the hypertrophic response and a direct progression to a cardiomyopathic phenotype. Significantly, this cardiomyopathic phase is associated with substantial cardiomyocyte death, which would indicate that ERK2 elicits a protective anti-apoptotic effect upon the cardiomyocyte (Ulm et al., 2014). Surprisingly, conditional deletion of both ERK alleles did not prevent pathological hypertrophic growth (Kehat et al., 2011). Further analysis revealed a selective role of ERK1/2 in the different forms of hypertrophy - while ERK1/2 mediates concentric growth responses to stimulus, as evidenced by dilation and decreased function of ERK1/2 KO mice as well as induction of fetal genes,, it prevents eccentric growth (Bueno et al., 2000; Kehat et al., 2011; Purcell et al., 2007). Surprisingly, cardiomyocyte-specific loss of *Erk2* in mice did not affect physiological cardiac remodelling in response to 4 weeks of swimming training, indicating independent pathways for adaptive hypertrophy in response to pathological or physiological stimuli (Ulm et al., 2014).

These diverse data from these different experimental models are perhaps not surprising given the complex regulation of this pathway, involving feedback at multiple levels that act to prevent constitutive activity. Together, these experimental findings suggest that ERK signalling contributes to acute, adaptive hypertrophic growth while repressing maladaptive growth thereby protecting the heart from maladaptive remodelling and progression to failure. A similar role is also suggested of certain AP-1 components where in their absence, the hypertrophic response is less adaptive in nature (Windak et al., 2013).

### MSK in the heart

As indicated above, ERK1/2 signalling is critical for transduction of hypertrophic cues to activation of IEG expression and induction of hypertrophy. Moreover, ERK1/2 phosphorylates hypertrophy-related transcription factors to modulate the activation. Phosphorylation of histone H3 at S10 and S28 has also been described as key events in the pathway indissociably linking MAPK activation and induction of IEG expression in response to mitogenic stimulation (Clayton and Mahadevan, 2003; Clayton et al., 2000). MAPK cannot however directly phosphorylate H3S28 and S10 but act via an intermediate kinase, which in fibroblastic cells was identified as the mitogen and stress activated kinase MSK1/2. MSK1/2 are promiscuous nuclear serine/threonine proteins. MSK is activated either through ERK1/2 or p38 MAPK cascades, resulting in phosphorylation of the MSK C-terminal Ca^2+/^calmodulin-dependent protein kinase (CaMK)-like domain, leading to a positive feed forward autophosphorylation event at the N-terminal AGC-like kinase domain (Roux and Blenis, 2004). The phosphorylated activated N-terminal domain then phosphorylates other substrates, including histone H3. Our study provides the first evidence for a role for MSK in promoter histone H3 phosphorylation and IEG induction in cardiomyocytes. Although MSK was identified as the kinase responsible for phosphorylation of histone H3 and which was required for robust IEG activation in cultured fibroblasts (Soloaga et al., 2003), the data presented here are the first demonstration that phosphorylation of H3S28 is mediated by MSK in response to pathological stressor in cardiomyocytes, and further that this event is required for IEG activation and mounting of the cardiomyocyte hypertrophic response. The specific role for MSK1/2 in responding to stress stimuli is consistent with the lack of overt phenotype in *Msk1/2* double KO mice (Wiggin et al., 2002b).

Histone H3 Ser 28 phosphorylation at IEG promoters is an important step in the induction of their expression. This phosphorylation event results in increased recruitment of BRG1, a component of the SWI/SNF remodelling complex to chromatin (Hang et al., 2010). Appropriate localisation of this complex to chromatin is likely a key step in bringing about the required transcriptional response. Notably, BRG1 is an important mediator of chromatin remodelling and transcriptional responses in cardiac development and disease. Indeed, BRG1 together with HDAC and PARP is recruited to the *Myh6/7* locus and is involved in bringing about the isoform switching of myosin heavy chain during cardiac maturation and in response to stress. The lack of BRG1 recruitment following DN-MSK expression described here would support these previous observations. Based on our data, we propose that H3S28 phosphorylation by MSK is an initial step in mediating this hypertrophic response. The recruitment of BRG1 further contributes to modulation of the epigenetic landscape of the heart through recruiting factors including EZH2 to acetylate H3K27 at enhancers of the mesoderm and for Polycomb-mediated repression of non-mesodermal genes (Gehani et al., 2010). Such a role for MSK-mediated phosphorylation of H3S28 is described in neuronal differentiation, where MSK1/2 phosphorylates and targets H3S28 in promoters with Polycomb repressor complex 2 (PRC2) – bound methylated H3K27, thereby displacing PRC2 for gene activation (Gehani et al., 2010). H3K27me3 is also lost at activated gene promoters in hypertrophy and disease in cardiomyocytes (Gilsbach et al., 2014; Thienpont et al., 2017). Whether loss of this mark is associated with gain of H3S28 phosphorylation is not determined in cardiomyocyte transcriptional responses, although elsewhere, phosphorylation of H3S28 has been shown to displace the PRC complex allowing H3K27 acetylation (Kim et al., 2012). Together, these data suggest that MSK-mediated phosphorylation of H3S28 is important in bringing about epigenetic changes underlying cardiac development and in the responses to hypertrophic cues.

Phosphorylation of histone H3 has previously been described in cardiomyocytes, albeit in the context of a bona fide hypertrophic response. In chronic remodelling in response to sustained sympathetic activation, CaMKII was found to bind directly and phosphorylate H3S28. In end-stage heart failure as well as in a murine model, CaMKII-mediated phosphorylation of H3S28 in the haemoglobin promoter results in enhanced expression in adult cardiomyocytes (Saadatmand et al., 2019). Whether this particular mechanism is protective or contributes to the pathological phenotype is undetermined. Importantly, CaMKII was found necessary for sustained elevation of global pH3S28, with major differences compared with control seen after 24 h catecholaminergic stimulation (Saadatmand et al., 2019). While CaMKII may indeed play a role at certain gene loci, the decrease in H3S28 phosphorylation in *Msk* KO animals, suggests that MSKs make an important contribution to maintaining phosphorylation of H3S28, particularly at IEG loci shown here, during disease remodelling. CaMKII is also shown to phosphorylate H3S10 leading to the induction of cardiac foetal genes (Awad et al., 2013). Notably, in the latter study, no CaMKII dependent phosphorylation of H3S28 was detected. The nuclear localisation of this kinase together with its identified role in HDAC phosphorylation provides a mechanism to remodel chromatin in a manner optimal for stimulation of MEF2-dependent gene expression during hypertrophic remodelling (Awad et al., 2013; Backs et al., 2006). CaMKII-associated pH3S10 is also sustained in end-stage heart failure (Awad et al., 2015). As we did not detect a robust change in H3S10 phosphorylation in NRVMs exposed to hypertrophic stimuli, we did not examine phosphorylation of this residue in disease in vivo. Together, CaMKII phosphorylation of H3S10 and H3S28 and H3S28 phosphorylation by MSK represent a potential mechanism that permits different stimuli at different phases of their action to selectively control the expression of discrete panels of target genes.

Consistent with studies elsewhere, we show that MSK lies downstream of ERK1/2, requiring ERK activity for function (Markou and Lazou, 2002) (Figure SB). MSK is also modified by p38 MAPK, which has been proposed to be required in addition for ERK1/2 for activation and in mediating pathological cardiac stress responses. Further, MSK is proposed to induce hypertrophic responses via CREB in a manner that also requires PKA (Markou et al., 2004). These studies were however constrained by the poor pharmacology of MSK with drugs used targeting PKC and PKA in the nM range (Alessi, 1997; Naqvi et al., 2012). Since in agreement with other studies, ERK inhibition was sufficient to prevent MSK activation and its regulation of effectors, we did not probe further the role of p38. Notably, as well as showing increased phosphorylation of sites indicative of its activation, *MSK1/2* gene expression was induced in response to hypertrophic stimulation both in vitro and in vivo. *MSK1/2* expression was also maintained in more chronic situations, further underlining the importance of this kinase in early responses to stress as well as in potentially sustaining it function. The persistence of IEG expression in the 6 week rat model of hypertrophic remodelling and in human disease may support this notion.

Other targets of MSK involved in the cardiac hypertrophic response have been described. The first substrate of MSK identified was the cAMP-Responsive Element-Binding Protein (CREB) transcription factor, which binds cAMP response DNA elements (CRE), associating with the histone acetyltransferase CREB-binding protein (CBP/P300) to activate transcription. CREB itself is also phosphorylated by a number of different kinases, including protein kinase A (PKA) (Johannessen et al., 2004). The role of MSK-activated CREB in vivo is controversial. Several studies have demonstrated that PKA-but not MSK-mediated CREB phosphorylation leads to CBP or p300 recruitment (Kasper et al., 2011). Cardiac-specific expression of a dominant negative form of CREB (DN-CREB) leads to a dilated cardiomyopathy phenotype (Watson et al., 2010). Given the extreme phenotype of DN-CREB in contrast to the relatively benign *Msk1/2* double KO, it is likely that normal CREB activity is independent of MSK in the heart. Related to its role in the nucleosomal response, MSK phosphorylates high mobility group 14 protein (HMG-14). This protein associates with phosphorylated H3 at activated promoters (Soloaga et al., 2003). Its role in chromatin remodelling remains elusive and its relationship with MSK and H3 phosphorylation remains however, to be determined (Phair and Misteli, 2000).

### Pharmacology of MSK for therapeutic targeting

The lack of a significant cardiac phenotype of *Msk1/2* KO that we describe suggests a limited role of MSK in the basal activity of cardiomyocytes. The stress-specific function of MSK may endow it however with possessing the necessary qualities of being therapeutically targetable. To date, no highly specific drugs suitable for in vivo use are available and those that are show efficacy at multiple other kinases important in cardiac remodelling, the use of which would preclude identification of MSK mechanisms of action and role in the heart (Naqvi et al., 2012). Specific inhibitors of kinases upstream of MSK, including members of the ERK/MAPK pathway have also been employed. The role of these kinases in the baseline activity of the heart is more significant making them less ideal targets for therapy.

## Conclusion

Our data identified MSK as the missing link between ERK/MAPK activation, histone H3S28 phosphorylation, IEG induction and cardiomyocyte hypertrophy induction. Further studies will lead to the identification of the wider significance of MSK-induced pH3S28 in the hypertrophic response and how it may be manipulated for therapeutic benefit.

## Materials and Methods

### Reagents

Chemicals purchased were from Sigma Aldrich and molecular biology reagents were from Thermo Scientific and Life Technologies, unless stated otherwise. Tables of antibodies and primers used in this study are included in Supplementary Tables and 3 and 4 respectively.

### Animal experiments

All experiments involving animals were in accordance with the European Directive 2010/63/EU. Experiments were performed in accordance with the UK Home Office and institutional guidelines or were approved by the Ethical Committee for Animal Experiments of the KU Leuven (Belgium). The Msk1/2 null animals have been previously described (Arthur and Cohen, 2000; Wiggin et al., 2002b). Hypertrophic remodelling was induced in 8-10 week old male mice animals by administration of isoproterenol (Iso, Sigma-Aldrich) at 10 mg/kg/day for one week via osmotic mini-pumps (Alzet) implantation as previously described and under project license P3A97F3D1 (Liu et al., 2009). Jugular vein infusion of ET-1 and Iso was performed as previously described (Archer et al., 2017) using an approved experimental protocol (license number P055/2017) approved by the Ethical Committee for Animal Experiments of the KU Leuven (Belgium), explained in detail later. Male Sprague Dawley rats subjected to six weeks of ascending aortic banding or a six-week treadmill training program were previously used to generate cardiomyocyte-specific nuclear (PCM-1 positive) RNA-sequencing data as previously described (Thienpont et al., 2017). RNA sequencing data was re-purposed for this study, focusing on the panel of IEGs. The sequencing data are available in the NCBI’s Gene Expression Omnibus (GEO) database (GEO GSE66653). Animals were housed and treated according to the European Directive 2010/63/EU.

### Echocardiography

Mice were anesthetised with Avertin (200 mg/kg). Cardiac function was assessed by transthoracic 2D M-mode echocardiography using an Acuson Sequoia C256 ultrasound system (Siemens) as previously described (Liu et al., 2009).

### Preparation of neonatal rat ventricular myocytes (NRVMs)

Primary neonatal rat ventricular myocytes (NRVMs) were isolated from 3-4 day old male and female Wistar pups and cultured as described previously (Higazi et al., 2009). Cultures were > 95 % pure. NRVMs were seeded at a density at which they exhibited spontaneous and synchronous beating throughout the experiment. 48 h after seeding, NRVMs were washed into serum-free medium (DMEM/M199 4:1, 1 mM sodium pyruvate, 5.5 µg/mL transferrin, 5 ng/mL sodium selenite, 1 X Antibiotic-Antimycotic (Life Technologies), and 3 µM cytosine b-D-arabinofuranoside (araC) and serum-starved for 24 h. NRVMs were subsequently stimulated with the agents described. Adenoviral infections were performed by incubation with a volume of virus-containing serum-free medium sufficient to cover the cells for 4 h. Agonist treatments diluted in serum-free medium were applied 24 h post-infection with adenovirus. Endothelin-1 (ET-1, Millipore), isoproterenol hydrochloride (Iso, Sigma Aldrich) and PD184352 (PD, Sigma Aldrich) treatments were performed at a final concentration of 100 nM, 10 nM and 1 µM respectively. All cellular treatments with PD were pre-treated with PD for 30 min prior to hypertrophic agonist application (ET-1/Iso). Control cellular experiments (no treatment) were treated with the same volume of vehicle only (DMSO for ET-1 and PD).

### Isolation and culture of adult rat ventricular myocytes (ARVMs)

Male Wistar rats (Harlan; ∼200 g) were anaesthetised by CO_2_ inhalation and killed by cervical dislocation. ARVMs were isolated by collagenase digestion following Langendorff perfusion as previously described and cultured in adult cardiomyocyte medium (M199, 1 % penicillin-streptomycin-l-glutamine, 0.2 % bovine serum albumin (BSA)) on laminin-coated (25 µg/ml) dishes (Drawnel et al., 2012). Adenoviral infections were performed for 12 h in a minimal volume of virus-containing medium. For experiments involving acute stimulation with ET-1 and Iso, cells in Tyrode were plated onto laminin-coated 8 well Nunc cover glasses cells, allowed to attach for 1 h at 37 °C, after which the Tyrode solution was replaced with Tyrode containing DMSO vehicle or Tyrode containing 10 nM PD. After 20 min, buffer was exchanged for Tyrode containing 100 nM ET-1 or 10 nM Iso ± PD. After 15 min, dishes were placed on ice and processed for immunostaining and imaging.

### Isolation of human ventricular cardiomyocytes

Donor human tissue was collected under a study protocol approved by the ethical committee of UZ Leuven (S58824), conformed to the Helsinki declaration, and was con-ducted in accordance with the prevailing national and European Union regulations on the use of human tissues. Donor information is displayed in Table S1.

Human ventricular myocytes were prepared from the explanted hearts immediately after removal as previously described (Dries et al., 2018). The explants hearts were collected in ice-cold modified Tyrode’s solution at the time of surgery (in mM: NaCl 130, KCl 27, HEPES 6, MgSO_4_ 1.2, KH_2_PO_4_ 1.2, glucose 50; pH 7.2 with NaOH) and transported to the lab. A wedge of the left ventricle with its perfusing coronary artery was cannulated. If possible, a wedge from the left anterior descending artery was cannulated, otherwise a left circumflex branch was used. The artery and the tissue was perfused at 37 °C with a Ca^2+-^free Tyrode’s solution (in mM: NaCl 130, KCl 5.4, HEPES 6, MgSO_4_ 1.2, KH_2_PO_4_ 1.2, glucose 20; pH 7.2 with NaOH) for 30 min followed by enzyme perfusion for 40 min (collagenase A, Roche and protease XIV, Sigma Aldrich in Ca^2+-^free solution) and after digestion, perfused with low Ca^2+^ Tyrode (Ca^2+-^free solution with 0.18 mM CaCl_2_ added). The digested tissue was minced, the suspension was filtered and the isolated myocytes were resuspended in normal Tyrode. After isolation, the cells were allowed to recover for 1 h before starting experiments or fixation. For experiments involving stimulation with Iso or ET-1, cells were plated onto poly-L-lysine coated 8 well Nunc cover glasses and allowed to attach for 1 h at 37 °C. After this period the Tyrode solution was replaced with Tyrode containing DMSO vehicle or Tyrode containing 1 µM PD. After 20 min, buffer was exchanged for Tyrode containing 100 nM ET-1 or 10 nM Iso ± PD. After 15 min, dishes were placed on ice and processed for immunostaining and imaging.

### Isolation of human cardiomyocyte nuclei

Nuclei from post-mortem left ventricular tissue were isolated and flow sorted according to pericentriolar material 1 (PCM-1) staining as previously described (Bergmann and Jovinge, 2012). 500,000 nuclei were sorted into 1 ml TRIzol reagent for RNase inhibition prior to RNA isolation. Human LV samples were obtained from the KI Donatum, Karolinska Institutet, Stockholm, Sweden, with permission for the analysis of human tissue for research purposes granted by the Regional Ethics Committee in Stockholm, Sweden. Donor information from which LV cardiomyocyte nuclei were isolated is displayed in Table S2.

### Histology and section preparation

Adult hearts were dissected from *Msk1/2* KO mice and covered in a layer of Tissue-Tek optimum cutting temperature (OCT) in Tissue-Tek® Cryomold® Molds (15×15×15 mm) and flash-frozen in liquid nitrogen-cooled isopentane (VWR). 10 μm ventricular sections were cut using a Leica cryostat and attached to SuperFrost Plus™ slides (VWR). Slides were frozen at -20°C prior to immunostaining.

Sections were thawed and rehydrated in phosphate buffered saline (PBS) for 5 min, followed by 15 min fixation in 4 % paraformaldehyde (PFA). After three washes (5 min each) in PBS, sections were permeabilised for 30 min in 0.2 % Triton X-100 in PBS (PBS-TX), then washed twice with PBS. Non-specific protein binding sites were blocked by incubation in PBS-TX containing 3 % BSA or 5 % goat serum for 1 h. Primary antibodies were diluted in blocking buffer and incubated overnight at 4 °C. After overnight incubation, slides were washed X3 in PBS and secondary antibodies added in blocking buffer for 1 h at room temperature. After incubation, the slides were washed and mounted in VECTASHIELD Antifade Mounting Medium containing DAPI (Vectorlabs). Confocal images were acquired using a Nikon A1R confocal microscope using a 40X 1.3 Numerical Aperture (N.A.) oil immersion objective.

### Picro Sirius Red staining for fibrosis analysis

10 µm thick sections were cut from OCT embedded tissue as above. Subsequently, sections were rehydrated and stained for collagen using a Picro Sirius red staining kit (PolySciences). After staining, sections were mounted in dibutylphthalate polystyrene xylene mounting medium. Images were acquired using a Zeiss Axioplan microscope configured with an Axiocam HrC camera. Polarization microscopy was performed on the Sirius red stained sections to visualize collagen type I and III based on the birefringence properties of collagen. The degree of fibrosis was quantified using Axiovision analysis software.

### Immunofluorescence analysis

Analysis of surface area of NRVM was carried out as previously described (6). Briefly, NRVMs for immunofluorescence were cultured and fixed in black 96-well imaging microplates (BD Biosciences). NRVMs were immunostained with primary antibodies against α-Act and ANF and detected using Alexa Fluor 488 and 568-coupled secondary antibodies (Table S3). After immunostaining, nuclei were labelled with Hoescht (1 µg/ml in PBS for 20 min). Images were captured using a BD Pathway 855 high-content imaging system and Attovision software. Cell planimetry was performed using ImageJ by drawing around the edge of the cells (NIH). At least 400 cells from three independent experiments were analysed. These images were also used for quantification of ANF protein expression as determined by counting the number of NRVM exhibiting a peri-nuclear ring of ANF.

For confocal imaging, NRVMs were cultured in 16-well chamber slides (Nunc). Slides were mounted onto a coverslip using VECTASHIELD Mounting Medium containing DAPI and sealed with clear nail varnish.

For staining of ARVMs and human isolated cardiomyocytes, cells were fixed and permeabilised in ice-cold 100 % methanol and incubated at -20 °C for 10 min. Methanol was washed from the coverslips twice with PBS and further permeabilisation performed by the addition of 0.5 ml ice-cold 100 % acetone and incubation at -20 °C for 1 min. Following an additional two washes in PBS, antibody labelling was performed as described for NRVM. To further reduce background staining, blocking and secondary antibody buffers contained 1 % BSA in addition to goat serum.

Immunofluorescence analysis of cardiac sections was performed as previously described (8). Snap-frozen heart samples were embedded in OCT (VWR), cryo-sections (thickness 10 µm) were collected on SUPERFROST PLUS microscope slides (VWR), fixed in 4 % PFA in PBS for 15 min, permeabilised in 0.25 % Triton-X100 in PBS for 15 min, and blocked in 5 % Chemibloc in PBS with 0.1 % Triton-X100 (PBS-TX) for 1 h. Sections were subsequently incubated overnight at 4°C in blocking buffer with primary antibodies as per Table S3. Samples were washed extensively in PBS-TX, and incubated with Alexa Fluor® secondary antibodies (Invitrogen) at 1:500 in PBS-TX for 1 h at room temperature. Where required, DAPI was included to identify the DNA in nuclei, respectively.

Sections were mounted in VECTASHIELD with DAPI (Vector Labs) and imaged on a Nikon A1R confocal microscope through a Plan Fluor DIC H N 40x oil immersion objective (N.A.=1.3). Image stacks were collected over a 2 µm stack thickness (0.2 µm z-step). Image stacks were analysed with Volocity Image analysis software (version 6.2.1, Perkin Elmer). Cardiac myocyte nuclei were identified by PCM-1- or Nesprin-positive labelling, as previously described (Thienpont et al., 2017).

Cellular apoptosis was measured in *Msk* KO mouse cardiac sections using the TACS® 2 TdT-DAB In Situ Apoptosis Detection Kit (Bio-Techne Ltd).

Confocal images were acquired using an Olympus FV1000 point scanning microscope attached to an Olympus IX81, equipped with a 40X/1.3 NA UPlanFI oil immersion objective or using a Nikon A1R confocal attached to a Nikon Ti microscope equipped with 40X 1.3 N.A. oil immersion objective.

### Histone isolation by acid extraction

NRVMs in 6-well dishes were washed once in ice-cold PBS. 0.5 ml of fresh ice-cold PBS was added to each well, the cells scraped and placed in pre-chilled 1.5 ml tubes. Cells were pelleted by centrifugation (10 min, 300 xg, 4 °C). PBS was removed and the pellet re-suspended in 1 ml hypotonic lysis buffer. The resuspended cells were incubated on a rotator at 4 °C, 30 rpm for 30 min. At the end of this incubation, intact nuclei were pelleted by centrifugation (10 min, 10000 xg, 4 °C). Nuclei were then resuspended in 400 µl 0.2 M H2SO4 by pipetting and vortexing. Histones were acid extracted overnight at 4 °C on a rotator at 30 rpm. Following acid extraction, nuclear debris was removed by centrifugation (10 min, 16000 xg, 4 °C) and the supernatant containing isolated histones transferred to a pre-chilled tube. To precipitate proteins, 100 % trichloroacetic acid was added to the supernatant in a drop-wise manner to achieve a final concentration of 25 %. The tube was gently inverted and then incubated on ice for 6 h. At the end of this period, precipitated proteins were recovered by centrifugation (10 min, 16000 xg, 4 °C). The supernatant was aspirated and acid removed from the tube by washing the pellet in 300 µl ice-cold acetone. After centrifugation (5 min, 16000 xg, 4 °C) and removal of the supernatant, the acetone wash and spin were repeated. Finally, the supernatant was gently removed and the pellet air-dried for 20 min at room temperature. The dried pellet was resuspended in 50 µl water and incubated overnight at 4 °C on a rotator at 30 rpm to maximise protein solubilisation.

### Immunoblot analysis

Cultures of NRVM were washed once in ice-cold PBS after which 80 µl pre-chilled RIPA buffer was added to the dish and incubated for 5 min on ice (25 mM Tris-HCl, pH 7.6, 150 mM NaCl, 0.1 % SDS, 1 % NP-40, 1 % Sodium Deoxycholate supplemented with 1 X Protease and Phosphatase inhibitor cocktails (Sigma Aldrich). The cell lysate was transferred to a pre-chilled tube and debris removed by centrifugation (5 min, 10 000 g, 4 °C). The supernatant was transferred to a clean tube and total protein concentration determined using the BCA assay (Thermo Scientific). Equivalent amounts of protein (10-30 µg) were loaded and samples prepared with LDS sample buffer (Invitrogen, final concentration 25 %) containing 2.5 % β-mercaptoethanol and boiled at 95 °C for 5 min before centrifuging briefly to remove debris.

Proteins were resolved on pre-cast 4-12 % NuPAGE 1.5 mm 10 well SDS gels (Invitrogen). The gels were rinsed with deionised water and placed in an X-cell Surelock Mini-cell running tank. The inner buffer chamber was filled with sufficient 1 X MOPS SDS Running buffer (Invitrogen) to cover the wells which were then rinsed with buffer expelled from a needle and syringe. 500 µl NuPAGE Antioxidant (Invitrogen) was added to the inner buffer chamber. The outer buffer chamber was filled with approximately 600 ml 1 X MOPS SDS Running buffer. 12 µl of Novex pre-stained sharp protein markers (Invitrogen) were loaded into the first well, followed by the boiled samples. Electrophoresis was performed at 200 V until the tracking dye reached the end of the gel.

For detection of ERK or MSK, proteins were transferred to a PVDF (0.45 µm, Millipore) membrane, which had been activated by immersion for 100 % methanol for 15 s and then placed in deionised water for 2 min. For detection of histone H3, proteins were transferred to a nitrocellulose (0.2 µm, Whatman). Proteins were detected with appropriate primary antibodies and HRP-conjugated secondary antibodies (Table S3). Immunoreactive bands were detected by enhanced chemiluminescence (Pierce).

### Reverse transcription quantitative PCR (RT-qPCR)

RNA was isolated from NRVMs and ARVMs using the RNeasy Micro Kit (Qiagen) and DNA removed by an on-column DNA digestion step. RNA was isolated from adult rat LV tissue, *Msk1/2* KO mouse LV tissue and human cardiomyocytes using TRIzol reagent (Invitrogen).

500-750 ng RNA was reverse transcribed using Superscript II (Invitrogen), the final cDNA synthesis reaction diluted 1/10 - 1/20 in nuclease free water and stored at -20 °C until required. Primer sequences were as previously described, unless otherwise indicated (Table S4), and were designed to span intron-exon boundaries to avoid amplification of genomic DNA (Higazi et al., 2009). The stability of a panel of reference genes was assessed using the GeNorm method for the experiments performed, and the most stable selected (Vandesompele et al., 2002). Three or four reference genes were selected for each set of experimental conditions for normalisation of gene expression qPCR analysis based on their stability for each set of samples and reaction conditions. Final primer concentration was 200 nM for all targets.

Reactions were performed on a LightCycler® 480 System (Roche) or on a CFX384 (BIO-RAD) in a 384-well format using Platinum SYBR Green qPCR SuperMix-UDG (Life Technologies). Expression analysis was carried out using the comparative ΔCt method as described (Livak and Schmittgen, 2001).

### Chromatin-immunoprecipitation (ChIP)

To NRVMs in 6-well dishes in culture medium, formaldehyde was added to a concentration of 1 % and incubated for 10 min on a rocking platform at room temperature. Cross-linking was terminated by the addition of 125 mM glycine for 10 min at room temperature. After washing once in ice-cold PBS, cells were collected into 0.5 ml PBS by scraping and subsequent centrifugation (5 min, 600 xg, 4 °C). Cell pellets were re-suspended in 1 ml ice-cold hypotonic membrane lysis buffer (50 mM Tris-HCl, pH 7.5, 5 mM EDTA, 140 mM NaCl, 1 % Triton X-100, 0.5 % NP-40 supplemented with 1 X Protease and Phosphatase inhibitor cocktails). The released nuclei were pelleted by centrifugation (3 min, 12000 xg, 4 °C) and then re-suspended in 200 µl SDS lysis buffer (50 mM Tris-HCl, pH 7.5, 10 mM EDTA, 1 % SDS supplemented with 1 X Protease and Phosphatase inhibitor cocktails). Cross-linked chromatin was fragmented by sonication using a pre-chilled Diagenode Bioruptor on the high power setting for three x 5 min cycles of 30 s ‘on’, 30 s ‘off’. The sonication protocol produced fragments predominantly below 500 bp. Wash buffer + (10 mM Tris-HCl, pH 7.5, 140 mM NaCl, 1 mM EDTA, 0.5 mM EGTA, 1 % Triton X-100, 0.1 % SDS and 0.1% Na deoxycholate supplemented with 1 X Protease and Phosphatase inhibitor cocktails) was then added to the lysate. After removal of remaining debris by centrifugation (10 min, 12000 xg, 4 °C), the supernatant (sonicated chromatin) was processed for immunoprecipitation.

Proteins of interest were precipitated using antibodies pre-conjugated to Dynabeads Protein A (Invitrogen). To this end, beads were washed in 4 changes of wash buffer and collected by the use of a DynaMag Magnet. Per ChIP, 10 µl washed beads and 5 µg Brg1 or phosphorylated H3S28 antibody were added to 90 µl wash buffer and incubated for two h at 40 rpm on a rotator at 4 °C. For the negative control ChIP, beads were used in the absence of specific antibody.

Prior to IP, wash buffer was removed from pre-prepared antibody-bead complexes and 200 µl chromatin added to each tube. 200 µl chromatin was reserved from each experimental condition as an Input sample. Chromatin was incubated with the antibody-bead complexes overnight at 4 °C on a 40 rpm rotator after which unbound chromatin was washed from the beads. Wash buffer was used for the first two washes, followed by one wash in high-salt wash buffer (wash buffer + with 500 mM NaCl) and finally two washes in TE buffer. At the end of the washes, the solution was transferred to a fresh tube and TE buffer removed from the beads. Elution of chromatin from the beads and protein digest with proteinase K were combined into one step. To this end, 150 µl complete elution buffer (20 mM Tris-HCl, pH 7.5, 50 mM NaCl, 5 mM EDTA, 1% SDS and 100 µg/ml proteinase K (Sigma)) was added and the beads incubated for 4 h at 68 °C with shaking at 1300 rpm. The supernatant was removed from the tube and 150 µl elution buffer (complete elution buffer without SDS and Proteinase K) added. After a further 5 min incubation at 68 °C, the two supernatants were combined. Input samples were processed in parallel with the ChIP samples. DNA was purified from each supernatant using the QIAEX II Gel extraction kit (Qiagen) following the manufacturer’s protocol for concentrating DNA from solutions. DNA was eluted in 40 µl buffer EB and stored at -20 °C until required.

Precipitated DNA for each experimental condition and antibody was quantified by qPCR using SYBR-GreenER in 12.5 µl reactions performed in triplicate (Invitrogen). Primers were designed to amplify the promoter sequences of c-Fos and c-Jun and the sequences of the primers used are given in appendix A. Primer binding sites were selected that encompassed predicted transcription factor binding sites. qPCR was performed using a CFX96 (BIO-RAD) or Roche Light Cycler480 real-time PCR instrument and cycling parameters were taken from the manufacturer’s instructions for SYBR-GreenER (Thermo Life Technologies).

The Ct values from triplicate technical replicates (n=3) from each sample were averaged to generate SampleCT and InputCt values. The values were analysed by expressing enrichment of the immunoprecipitated DNA for each antibody as a percentage of the input sample for the relevant experimental condition. ChIPs were repeated on at least 3 independent experimental samples.

### Jugular vein infusion of endothelin-1/isoproterenol in Wistar rat

Experimental protocols were approved by the local ethical committee (Ethische Commissie, Dierproeven, KU Leuven), under license number P055/2017. 250-300 g Wistar (RccHan:WIST) male rats were obtained from Harlan (NL). Anesthesia was induced using ketamine and xylazine in combination (100 mg/kg ketamine, 10 mg/kg xylazine) by IP. Body temperature was maintained throughout the procedure with a heated mat (Sanitas).

A small area of chest was shaved with depilatory cream (Veet) and limbs secured with tape. A small incision was made just above and to the right hand side of the sternum and the skin stretched thin with a hemostat to make the jugular vein visible. A 30 gauge needle attached to a cannula (2F x 30 cm, green, Portex) was inserted into the jugular vein, just before it branches and the vein disappears under the pectoral muscle.

The cannula was attached to a 5 ml syringe and a dispensing pump (Harvard Apparatus) dispensing the required volume (300-500 µl) over a 15-min period. A slow steady release of the dosage in this manner was required to reduce the acute vasoconstrictive effect of a single rapid injection of the same dosage.

Endothelin-1 (Millipore) was administered at a final dosage of 1000 ng/kg and isoproterenol hydrochloride (Sigma Aldrich) at 50 µg/kg. Final working concentrations prepared in sterile saline and vehicle-only controls (Ctrl) were administered the same volume of sterile saline over a 15 min period. On withdrawal of the needle, medical gauze was placed over the wound and pressure applied until bleeding stopped. The wound was cleaned with iodine solution and for the 24 h time point, the skin was sutured together with interrupted stiches.

For the 15 min time point, rats were sacrificed by cervical dislocation and heart removed for dissection immediately. For the 24 h time point, rats were allowed to recover alone in a cage on heated mat, with easy access to food and water. The humane 24 h end-point was performed by anesthesia induction in an isoflurane chamber followed by cervical dislocation and immediate removal of the heart.

Whole hearts were removed and placed in ice cold PBS briefly to remove excess blood, dissected using a sterile surgical scalpel in PBS on ice and weighed on a microbalance before snap-freezing in liquid nitrogen and stored at -80 °C.

### Adenoviral methods

Adenoviruses were produced and amplified in HEK293 cells and purified as previously described (Archer et al., 2017). Adenoviruses to express the WT and catalytically dead D565A mutant (DN) of MSK1 were generated using the AdEasy method by sub-cloning the cDNA for MSK1 or its mutant from a pCMV5 backbone (kindly provided by Prof D Alessi, University of Dundee) into pShuttle CMV (Alessi, 1997). PacI digested recombinant plasmids were transfected into HEK293 cells and crude adenovirus harvested after 10-14 days. Adenoviruses for dominant negative (DN)-Jun and AP-1 luciferase were purchased from Vector Biolabs. All viruses were amplified in HEK293 cells, purified using the Vivapure Adenopack 100 (Sartorius) and titrated by end-point dilution in HEK293 cells.

### Analysis of luciferase reporter activity

The AP-1 luciferase reporter was expressed using an adenoviral vector and luciferase activity determined using a luciferase assay kit from Promega as previously described (Higazi et al., 2009). Briefly, cultures of NRVM in 48 well plates were infected in duplicate and agonist treatments applied for 24 h. After removal of medium, cells were lysed in 150 µl 1X cell culture lysis buffer (Promega). Luciferase activity present in 10 µl of lysate clarified by centrifugation was then quantitated in white 96-well luminometer plate (Microlumat Plus 1b 96V instrument, Berthold Technologies) using 50 µl of luciferase assay reagent (Promega).

### Small interfering RNA (siRNA) knockdown

Stealth™ siRNAs were purchased from Invitrogen. To achieve sufficient knockdown of *Msk1* or *Brg1*, two siRNAs targeting different regions of the target mRNA were selected. Medium GC-content non-silencing siRNA (Invitrogen) was transfected as a negative control. Transfections were performed at the onset of the serum starvation period using Dharmafect I and Accell medium (Dharmacon). To transfect NRVM cultured in 12-well dishes, 200 pmol siRNA duplexes were made up to 100 µl total volume in Accell media and the solution mixed by pipetting. In a separate tube, 6 µl Dharmafect I was added to 94 µl Accell medium and mixed. After incubation for 5 min at room temperature, the tubes were combined, mixed and incubated for a further 20 min at room temperature. During this incubation, NRVM culture medium was replaced with 800 µl pre-warmed Accell medium. At the end of the 20 min incubation, transfection complexes were added to the cultures in a drop-wise manner and incubated with the cells for 6 h. Post-transfection, Accell medium was replaced with fresh maintenance medium and the remainder of the serum starvation period carried out.

### Statistical analysis

Data were collated in Microsoft Excel and statistical analysis performed using GraphPad Prism v7.0 or v8.0. Data is presented as the mean of at least three independent experiments ± the standard error of the mean (SEM). The number of independent experiments for each figure is indicated in the figure legend. For comparison between two groups, p-values were calculated using the unpaired two-tailed t-test. To calculate p-values for data comparing three or more groups, one-way ANOVA with Bonferroni’s multiple comparison test for p value correction. P-values less than 0.05 were taken as significant. Individual (adjusted) p values are indicated on the figures.

## Acknowledgments

We would like to thank Dario Alessi (FRS FRSE FMedSci, University of Dundee) for providing the pCMV5 backbone vector for sub-cloning of MSK1 and Peter Pokreisz (CoBioRes NV and KU Leuven) for assistance in establishing techniques for jugular vein infusion of rats.

## Author contributions

Conceptualisation Ideas; E.L.R., F.D., S.M., H.L.R.

Data curation and Formal analysis; E.L.R., F.D., S.M., C.R.A., W.L., H.O., K.A., J.M.A., C.K.N., S.C.A., H.L.R.

Investigation; E.L.R., F.D., S.M., C.A., H.O., K.A., J.M.A., C.K.N., W.L., H.L.R.

Writing - Original Draft Preparation; E.L.R., F.D., H.L.R.

Writing - Review & Editing; all authors

Resource provision; O.B., K.A., J.M.A., I.S., K.R.S.

Supervision Oversight and Leadership; S.C.A., I.S., K.R.S., O.B., H.L.R.

## Abbreviations

AB: Ascending aortic banding
ANF/Nppa: atrial natriuretic factor
BNP/Nppb: brain natriuretic factor
CVD: cardiovascular disease
ChIP: chromatin immunoprecipitation
ara-C: cytosine b-D-arabinofuranoside
CaMKII: Ca^2+^/calmodulin-dependent protein kinase
ET-1: endothelin-1
ERK1/2: extracellular signal-regulated kinase 1/2
GPCR: fractional shortening
FS: G-protein coupled receptor
HF: heart failure
H3: Histone 3
H3S10: Histone H3 Serine 10
p-H3S10: phosphorylated H3S10
H3S28: Histone H3 Serine 28
h: phosphorylated H3S28, pH3S28, hours
IEG: immediate early gene
KO: knockout
MAPK: mitogen activated protein kinase
MSK: mitogen and stress activated kinase
Iso: isoproterenol
min: minutes
α-MHC: myosin heavy chain, α isoform
β-MHC: myosin heavy chain, β isoform
NRVMs: neonatal rat ventricular myocytes
NFAT: nuclear factor of activated T cells
PD: PD184352 (inhibitor of the upstream kinase of ERK1/2, MEK1/2)
PWd: posterior wall thickness in diastole
SRF: serum response factor
TAC: transverse aortic constriction
siRNA: small interfering RNA
WT: wild type.

---

**Figure Supplement 1:**
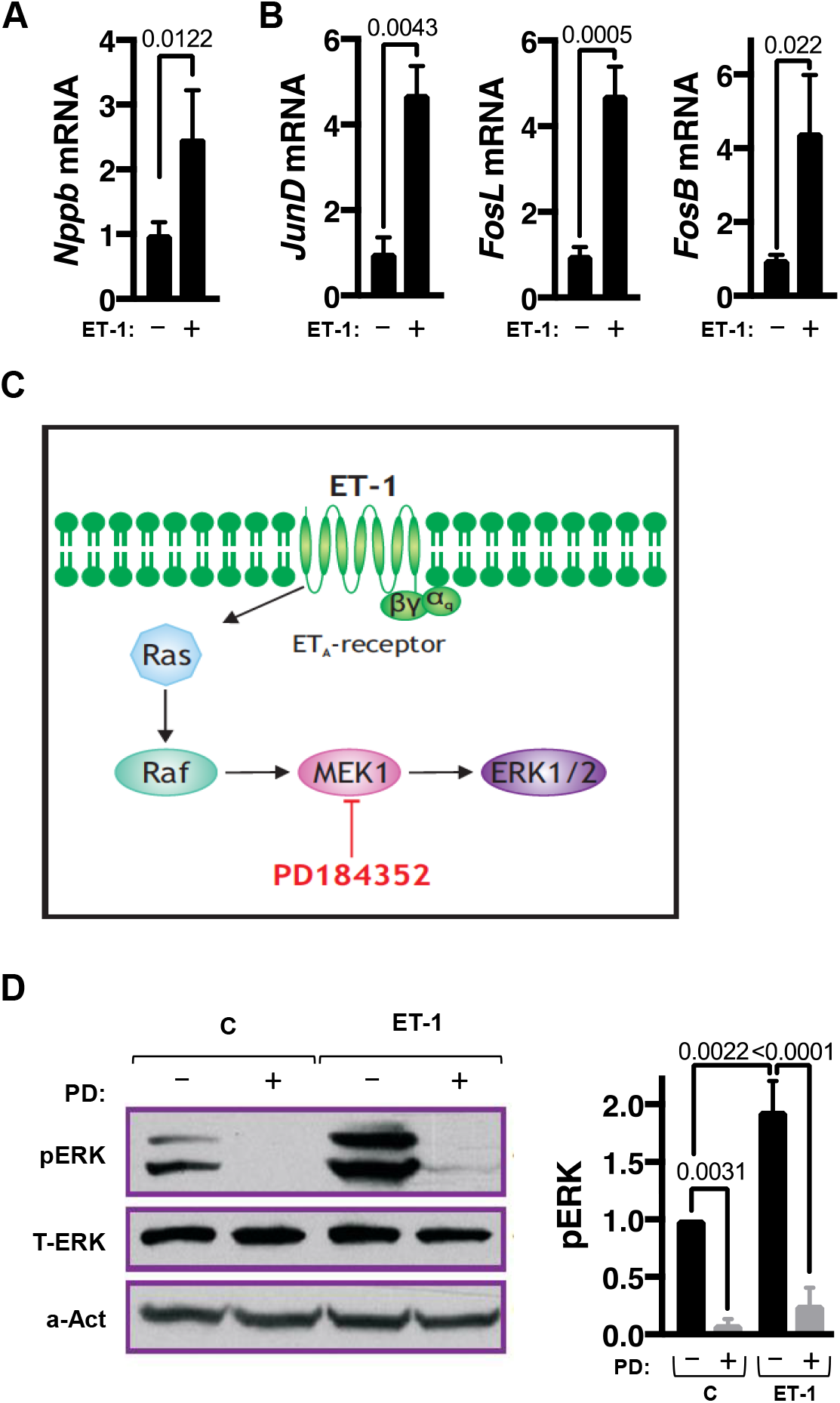
IEG induction is activated by acute hypertrophic stimulation and attenuated by suppression of ERK signalling in vitro. **A**. RT-qPCR analysis of *Nppb/Bnp* mRNA expression in NRVMs 24 h after ET-1 exposure. N=4, n=3. **B**. RT-qPCR gene expression analysis of *JunD, FosL* and *FosB* mRNA in NRVMs +/- ET-1 treatment for 15 min. N=4, n=3. **C**. Schematic diagram of the kinase signalling cascade induced by ET-1 stimulation resulting in ERK1/2 activation. Pharmacological inhibition of MEK1 by PD184352 (PD) is indicated. **D**. Immunoblot analysis for phosphorylated ERK in NRVMs +/- PD +/- ET-1. Left: Representative immunoblot for phosphorylated ERK (pERK), normalised to total ERK (T-ERK) and sarcomeric α-Act. Right: Quantitation of relative p-ERK normalised to T-ERK and α-Act. N=5.

**Figure Supplement 2:**
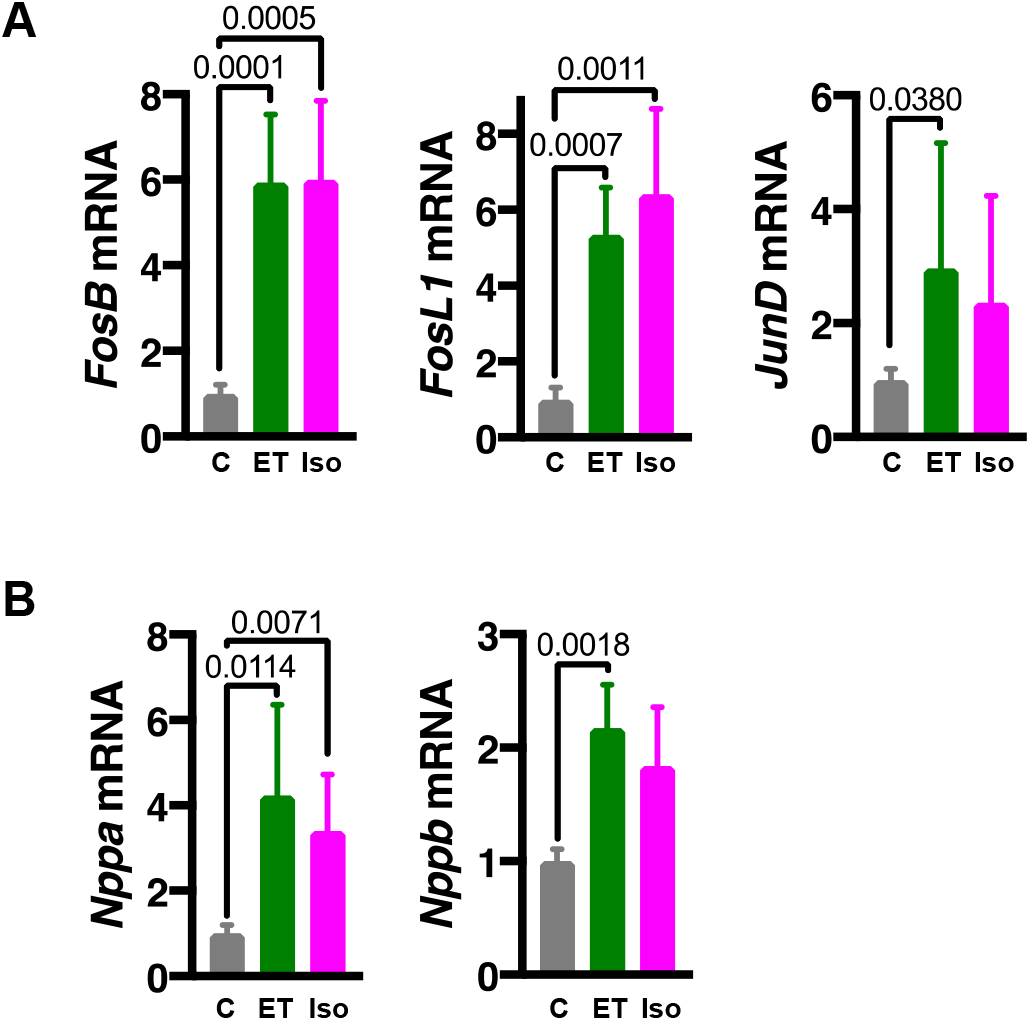
Histone H3S28 phosphorylation is associated with IEG induction and cardiac hypertrophy. **A**. RT-qPCR analysis of immediate early genes *JunD, FosL1, FosB* and of *Smarca4 (Brg1*) mRNA in hearts from adult male Wistar rats that were administered ET-1 or Iso through jugular vein administration and sacrificed 15 min later. C, N=7, ET-1, N=8 Iso, N=6, n=3. **B**. RT-qPCR expression analysis for cardiac hypertrophy-associated foetal genes *Nppa*/*Anf* and *Nppb*/*Bnp* mRNA in adult male Wistar rats that were administered ET-1 or Iso through jugular vein administration and sacrificed 15 min later. C, N=6, ET-1, N=6, Iso, N=7, n=3.

**Figure Supplement 3:**
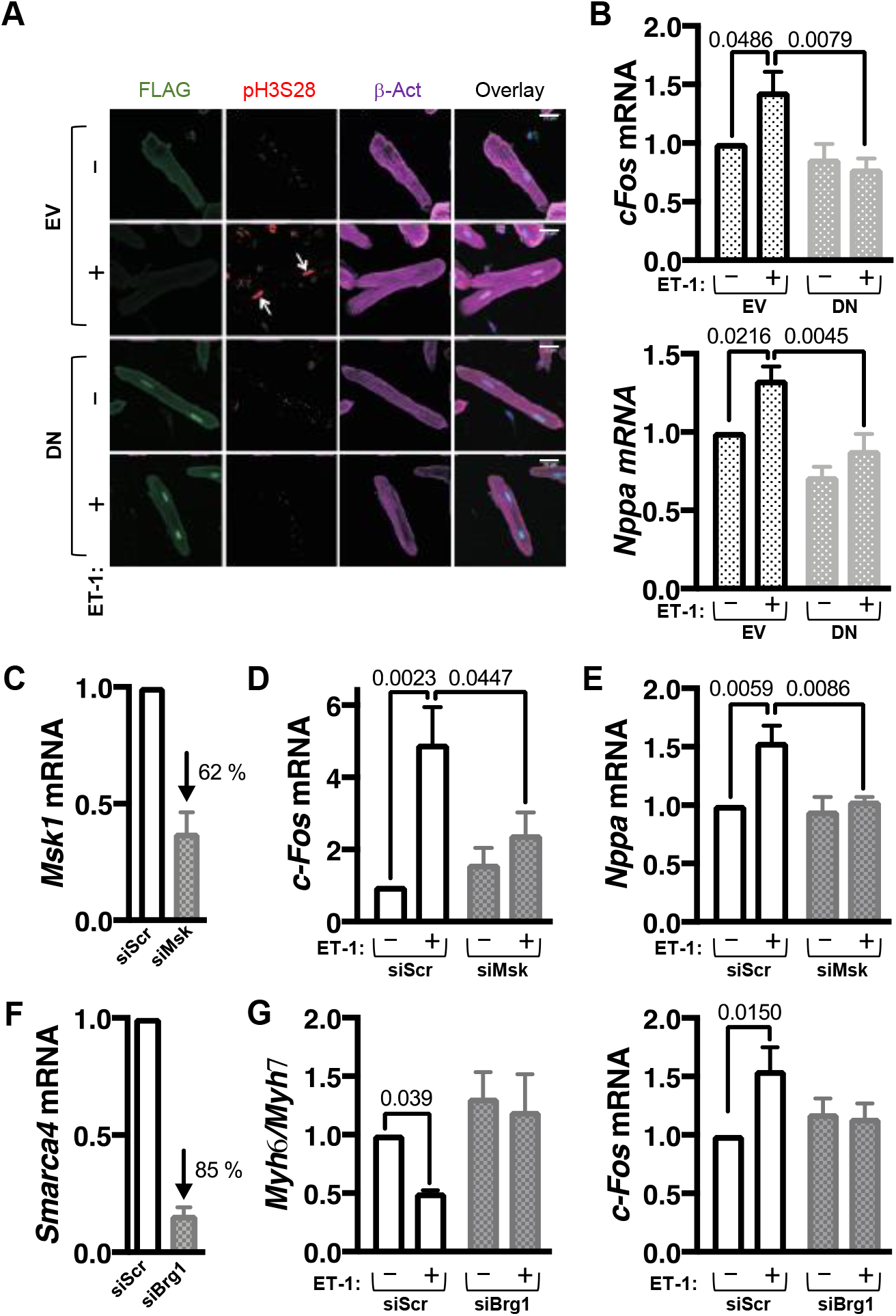
IEG induction is a conserved feature of hypertrophy in adult cardiomyocytes. **A**. Representative confocal images of immunostained ARVMs infected with adenovirus expressing FLAG-tagged DN-Msk (kinase dead). ARVMs are stained for FLAG (green), pH3S28 (red) and β-Act (purple). FLAG-tagged DN-Msk is enriched in the nuclei in ARVMs. **B**. RT-qPCR analysis of *c-Fos* (Top) and *Nppa*/Anf (Bottom) mRNA in ARVMs infected with EV or DN-MSK1 +/- ET-1 treatment for 15 min. For *c-Fos* mRNA (top), EV ctrl and EV ET-1, N=4, n=3. For DN-MSK ctrl and DN-MSK ET-1, N=3, n=3. For Nppa/Anf mRNA (bottom), N=3, n=3. Scale bar = 25 µm. **C**. Analysis of siRNA-mediated knockdown of *Msk1* in NRVMs. RT-qPCR analysis of *Msk1* mRNA expression in NRVMs transfected with scr or siMsk siRNA is shown. N=3, n=3. **D**. RT-qPCR analysis of *c-Fos* mRNA in NRVMs transfected with scr or siMsk siRNA +/- ET-1 for 15 min. N=4, n=3. **E**. RT-qPCR analysis of *Nppa/Anf* mRNA expression in NRVMs transfected with scr or siMsk siRNA +/- ET-1 for 15 min. N=3, n=3. **F**. Analysis of *Smarca4* knockdown in NRVMs. *Smarca4/Brg1* mRNA abundance was measured by RT-qPCR in NRVMs transfected with siRNA targeting Smarca3 (siBrg1) and compared with NRVMs transfected with scrambled control (scr) siRNA. N=3, n=3. **G**. RT-qPCR analysis of *Myh6/Myh7* (Left) and *c-Fos* (Right) mRNA expression in NRVMs transfected with scr or siBrg1 siRNA +/- ET-1 for 15 min. For *Myh6/Myh7* (Left), N=5. n=3. For *c-Fos* (Right), N=6, n=3.

**Figure Supplement 4:**
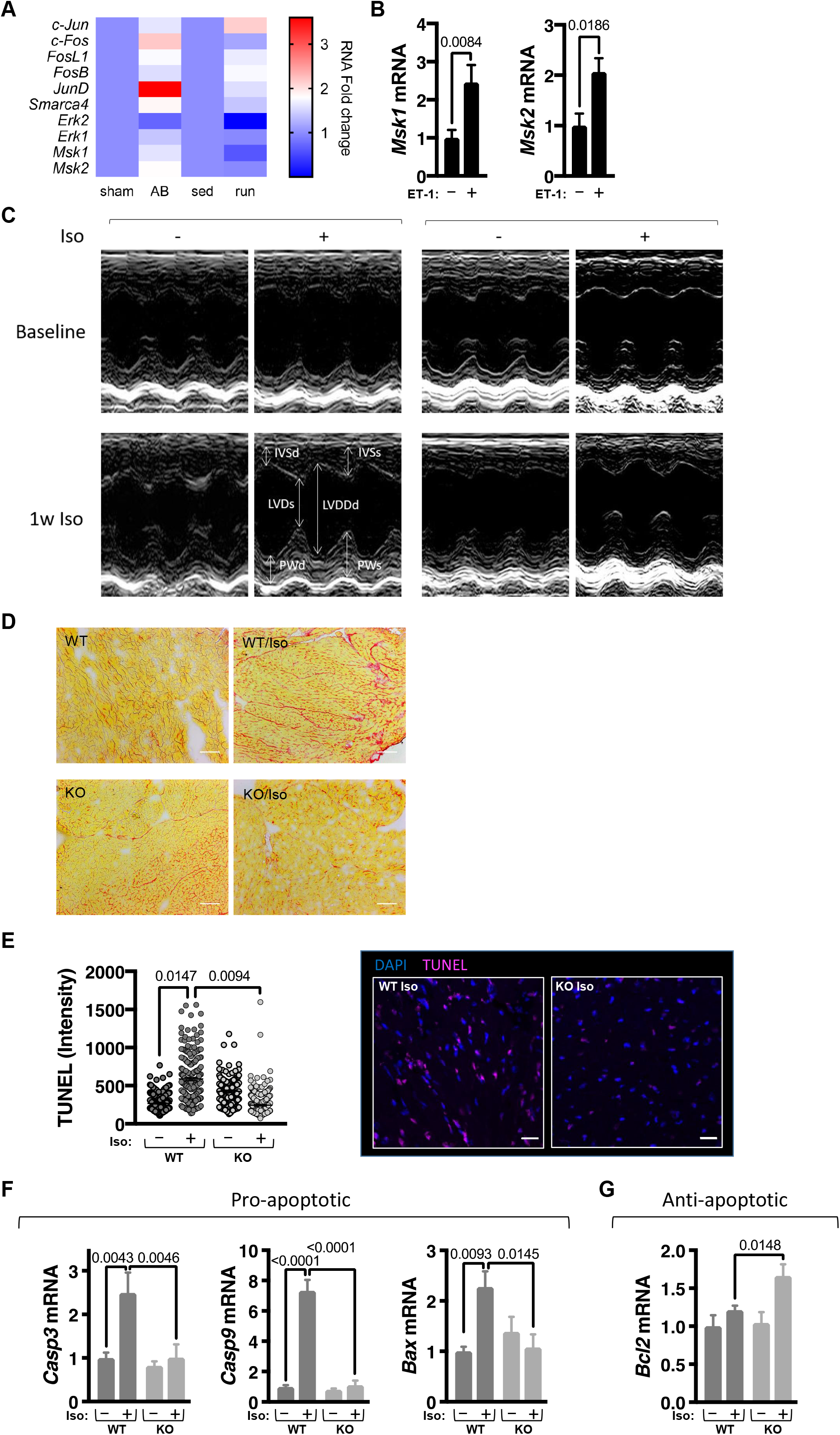
Genetic MSK inhibition attenuates IEG activation and cardiomyocyte hypertrophy in vivo. **A**. Heat map showing expression of immediate early gene (*c-Jun, c-Fos, FosL1, FosB, JunD*), *Smarca4/Brg1, Erk1/2* and *Msk1/2* in left ventricular Pcm-1+ve cardiomyocyte RNA-seq data in models of pathological (Ascending aortic banding, AB) or physiological (treadmill training, run) in male Sprague Dawley rats. Data extracted from (Thienpont et al., 2017). Data available at GEO GSE66653. **B**. RT-qPCR analysis of *Msk1* and *Msk2* mRNA expression in NRVMs treated with ET-1 for 24 h. N=4, n=3. **C**. Representative M-mode echocardiogram from WT/*Msk* KO mice +/- Iso for 1 week. Echocardiographic measurements indicated. IVSd=Interventricular septal dimension end systole, IVSd=Interventricular septal dimension end diastole, LVDs=Left ventricle diameter end systole, LVDd=Left ventricular diameter end diastole, PWd=posterior wall dimension end diastole, PWs=posterior wall dimension end systole. **D**. Representative left ventricular tissue sections from WT/*Msk* KO mice +/- Iso for 1 week showing Picro sirius red staining for collagen. The scale bar indicates 50 µm. **E**. Terminal deoxynucleotidyl transferase dUTP nick end labelling (TUNEL) staining of left ventricular cardiac sections in *Msk1/2* KO mice and wild type littermates +/- Iso infusion for 1 week. TUNEL assay measures fragmented DNA as a mark of apoptosis. Left: Quantification of TUNEL in cardiac nuclei. Right: Representative immunostaining images for TUNEL (green) and nuclei are stained with DAPI (blue). N=3, *<u>n</u>*=100-250. Scale bar indicates 20 µm. **F**. RT-qPCR analysis of pro-apoptotic marker genes caspase 3 (Casp3), caspase 9 (Casp9) mRNA expression in left ventricle from *Msk1/2* KO mice and wild type littermates +/- Iso infusion for 1 week. WT ctrl, N=5, WT Iso, N=4, KO ctrl, N=5, KO Iso, N=5, n=3. **G**. RT-qPCR analysis of the expression of the anti-apoptotic marker gene *Bcl2* mRNA expression in left ventricle from *Msk1/2* KO mice and wild type littermates +/- Iso infusion for 1 week. WT ctrl, N=5, WT Iso, N=4, KO ctrl, N=5, KO Iso, N=5, n=3.

