## Supplementary Tables for "MSK-mediated phosphorylation of Histone 3 Ser28 couples MAPK signaling with early gene induction and cardiac hypertrophy"

**Supplementary Data**

**Table S1: Human healthy donor samples for cardiomyocyte isolation**

| Patient | Age | Sex | BMI | Medications |
|---|---|---|---|---|
| HD-13 | 78 | F | 23 | Antidiuretics, Insulin, corticosteroids |
| HD-19 | 57 | F | 23 | Noradrenalin, Insulin, corticosteroids, T3 |
| HD-20 | 74 | F | 31 | Antibiotics, corticosteroids |
| HD-30 | 73 | F | 23 | Noradrenalin, Dopamine, Insulin, corticosteroids |

Samples used in Table S1 were used to generate data in Figures 5A-B.

**Table S2: Human control and hypertrophic left ventricular cardiomyocyte nuclei**

| Sample I.D. | Age (yrs) | B.M.I. | Heart Mass (g) | Pathology/Cause of Death |
|---|---|---|---|---|
| ND231 | 57 | 21.9 | 320 | Control/multiple injuries, fall down from 4th floor. |
| ND241 | 43 | 32.2 | 590 | left ventricular hypertrophy/natural |
| ND248 | 57 | 21.7 | 460 | Control/suicide |
| ND249 | 31 | 22 | 343 | Control/suicide |
| ND251 | 39 | 33.7 | 582 | Control/Intoxication |
| ND252 | 37 | 20 | 496 | left ventricular hypertrophy/natural |
| ND259 | 60 | 23.7 | 400 | Control/intoxication |
| ND279 | 61 | 29 | 710 | left ventricular hypertrophy/natural |
| ND288 | 63 | 30 | 563 | left ventricular hypertrophy/intoxication |

Samples in Table S2 were used to generate data in Figure 5C-F.

**Table S3: Antibody order information and concentrations**

IF = immunofluorescence, IB = immunoblotting

| Antibody | Order Information | Technique | Dilution Used |
|---|---|---|---|
| ANF | Bachem T-4014 | IF | 1:500 |
| pERK | Cell signalling technology (CST), 9106S | IB | 1:1000 |
| T-ERK | BD Biosciences, 610031 | IB | 1:5000 |
| α-Actinin | Sigma Adrich, A7811 | IB | 1:2000 |
| | | IF | 1:100 |

| | | | |
|---|---|---|---|
| B-Actin | Abcam, 52219 | IF | 1:1000 |
| Histone3 | Sigma Aldrich, H0164 | IB | 1:10 000 |
| Phos H3S10 | CST, 9701S | IB | 1:1000 |
| Phos H3S28 | Sigma Aldrich, H9908 | IB | 1:2000 |
| | | IF | 1:1000 |
| Phos H3S28 | Millipore, 07-145 | ChIP | 1:100 |
| PCM1 | Sigma Aldrich Prestige, HPA023370 | IF | 1:500 |
| Nesprin1 | MANNES1A (7A12) kindly donated to use by Glenn Morris) (1) | IF | 1:100 |
| pMSK | CST, 9591S | IB | 1:1000 |
| cFos | Abcam, ab190289 | IF | 1:250 |
| FLAG | Sigma Aldrich, FF3165 | IB | 1:2000 |
| | | IF | 1:100 |
| **Secondary Antibodies** | | | |
| HRP-conjugated antibodies | Jackson Laboratories | IB | 1:10 000 |
| Alexa 488 | Life Technologies, R37116, A-11029 | IF | 1:500 |
| Alexa 568 | Life Technologies, A-11036 | IF | 1:500 |
| Alexa 647 | Life Technologies, A-21244 | IF | 1:500 |

**Table S4: Primer sequence information**

Primers were ordered from Sigma Aldrich or Integrated DNA Techologies as lyophilized custom oligos, purified by desalting.

| Gene target | Species | Sequence |
|---|---|---|
| **RTqPCR** | | |
| GAPDH | Rat | F-CAAGATGGTGAAGGTCGGTGT<br>R-GGTCGTTGATGGCAACAATG |
| | Mouse | F-AATGGTGAAGGTCGGTGTGAAC<br>R-TCGTTGATGGCAACAATCTCC |
| | Human | F-GAGTCAACGGATTTGGTCGT<br>R-GACAAGCTTCCCGTTCTCAG |
| TBP | Rat | F-TGACTCCTGGAATTCCCATC<br>R-TGTGTGGGTTGCTGAGATGT |
| | Mouse | F-GGGGAGCTGTGATGTGAAGT<br>R-CAGGAGAACATGGCAGACAA |
| | Human | F-TATAATCCCAAGCGGTTTGC<br>R-GCTGGAAAACCCAACTTCTG |
| YWHAZ | Rat | F-ACCCACTCCGGACACAGAAT |

| | | |
|---|---|---|
| | | R- AGGCTGCCATGTCATCGTA |
| | Mouse | F-CAGAAGACGGAAGGTGCTGAGA<br>R-CTTTCTGGTTGCGAAGCATTGGG |
| | Human | F-CGAGATCCAGGGACAGAGTC<br>R-GGATGTTCTGTGTCCGGAGT |
| 18S | Rat/Mouse | F-ATCCATTGGAGGGCAAGTC<br>R-CGCTCCCAAGATCCAACTAC |
| | Human | F-AAACCACAGGCAAACACCTC<br>R-GCACTTTGGGTGGTCAAGTT |
| RPL32 | Rat/Mouse | F-GGCCAGATCCTGATGCCCAAC<br>R-CAGCTGTGCTGCTCTTTCTAC |
| | Human | F-AGGCATTGACAACAGGGTTC<br>R-GACGTTGTGGACCAGGAACT |
| SDHA | Rat | F-TCGCACTGTGCATAGAGGAC<br>R-ATGCCTGTAGGGTGGAACTG |
| | Mouse | F-AAGGCAAATGCTGGAGAAGA<br>R-TGGTTCTGCATCGACTTCTG |
| | Human | F-TCGCACTGTGCATAGAGGAC<br>R-GCCTGTAGGGTGGAACTGAA |
| LMNA | Rat | F-TGAGTACAACCTGCGCTCAC<br>R-TGTGACACTGGAGGCAGAAG |
| | Mouse | F-GCACCGCTCTCATCAACTCC<br>R-TCTTCTCCATCCTCGTCGTCA |
| | Human | F-CTACACCAGCCAACCCAGAT<br>R-GGTCGAAGGACAGAGACTGC |
| Tuba 6 | Mouse | F-GGCAGTGTTCGTAGACCTGGA<br>R-TTATTGGCAGCATCCTCCTTG |
| cFOS | Rat | F-CCGACTCCTTCTCCAGCATG<br>R-GTGGAGATGGCTGTCACCGT |
| | Mouse | F-GGGAATGGTGAAGACCGTGTCA<br>R-GCAGCCATCTTATTCCGTTCCC |
| | Human | F-GCCTCTCTTACTACCACTCACC<br>R-AGATGGCAGTGACCGTGGGAAT |
| cJUN | Rat | F-TTGAAAGCGCAAAACTCCGA<br>R-GTTAGCATGAGTTGGCACCC |
| | Mouse | F-CAGTCCAGCAATGGGCACATCA<br>R-GGAAGCGTGTTCTGGCTATGCA |
| | Human | F-CCTTGAAAGCTCAGAACTCGGAG<br>R-TGCTGCGTTAGCATGAGTTGGC |
| JUND | Rat | F-g a c a t g g a c a c g c a g g a a c<br>R-t c t g g c t t t t g a g g g t c t t g |
| | Human | F-ATCGACATGGACACGCAGGAGC<br>R-CTCCGTGTTCTGACTCTTGAGG |
| FOSB | Rat | F-a a a c a a a c a a a a c c g c a a g g<br>R-g g c g g t c a g a c a g a a g a g t c |
| | Human | F-TCTGTCTTCGGTGGACTCCTTC<br>R-GTTGCACAAGCCACTGGAGGTC |
| FOSL | Rat | F-a g a g c t g c a g a a g c a g a a g g<br>R-g c t g g t a c c a c c t g t g t c c t |

| | | |
|---|---|---|
| | Human | F-GGAGGAAGGAACTGACCGACTT<br>R-CTCTAGGCGCTCCTTCTGCTTC |
| SMARCA4 | Rat | F-CTACAGGAGCGGGAGTACAG<br>R-TGGTTGCTTTGGTTCGAAGG |
| | Mouse | F-GAAAGTGGCTCTGAAGAGGAGG<br>R-TCCACCTCAGAGACATCATCGC |
| | Human | F-CAAAGACAAGCACATCCTCGCC<br>R-GCCACATAGTGCGTGTTGAGCA |
| MSK1 | Rat | F-TCTATGTTGGAGAGATCGTGCTTG<br>R-AATCTGTCAGCACCACATGGC |
| | Mouse | F-TGGTCCATAGCACCTCTCAGCT<br>R-CTCTCCGCCATTCAGAAGTTCC |
| | Human | F-CCTGGAACACATTAGGCAGTCG<br>R-CACCTCATGCTCTGTGAAACGC |
| MSK2 | Rat | F-ATCCTGGACTATGTGAGCGG<br>R-CTAGGGCCAGCACAATCTCT |
| | Mouse | F-CAACGTGGTGAATCTGCATGAGG<br>R-AGCCGCTTCTTGCGGATGTGTT |
| | Human | F-TGTGGGCAACTTTGCGGAGGAA<br>R-GAGAATGGAGGGTGCCACAAAG |
| MYH6 | Rat | F-ACAGAGTGCTTCGTGCCTGAT<br>R-CGAATTTCGGAGGGTTCTGC |
| | Mouse | F-CAGAGGAGAAGGCTGGTGTC<br>R-CTGCCCCTTGGTGACATACT |
| | Human | F-CTGGGCAAGTCCAACAATTT<br>R-CCAGCCCAGGATGTTGTAGT |
| MYH7 | Rat | F-GCCTACAAGCGCCAGGCT<br>R-CATCCTTAGGGTTGGGTAGCA |
| | Mouse | F-CTTCAACCACCACATGTTCG<br>R-TCTCGATGAGGTCAATGCAG |
| | Human | F-GTGAAAGTGGGCAATGAGT<br>R-TGGTGAAGTTGATGCAGAGC |
| Nppa/Anf | Rat | F-CGTATACAGTGCGGTGTCCAAC<br>R-CCTCATCTTCTACCGGCATC |
| | Mouse | F-GTGCGGTGTCCAACACAGAT<br>R-TTCCTCAGTCTGCTCACTCAGG |
| Nppb/Bnp | Rat | F-GGTCTCAAGACAGCGCCTTCC<br>R-CTTCCTAAAACAACCTCAGCCCGTC |
| | Mouse | F-GCCAGTCTCCAGAGCAATTC<br>R-GTTCTTTTGTGAGGCCTTGG |
| COL1A1 | Mouse | F-GCCAAGAAGACATCCCTGAA<br>R-GCCATTGTGGCAGATACAGA |
| Casp3 | Mouse | F- GGGCCTGTTGAACTGAAAAA<br>R- CCGTCCTTTGAATTTCTCCA |
| Casp9 | Mouse | F- GCCAGAGGTTCTCAGACCAG<br>R- AAGCCGTGACCATTTTCTTG |
| Bax | Mouse | F- tgcagaggatgattgctgac<br>R- gatcagctcgggcactttag |
| Bcl-2 | Mouse | F- ggacttgaagtgccattggt |

| | | |
|---|---|---|
| | | R- a g c c c c t c t g t g a c a g c t t a |
| **ChIP-qPCR** | | |
| cFos promoter | Rat | F-CCTTGCGCTGCACCCTCAGA<br>R-CGGCCGTGGAAACCTGCTGA |
| cJun promoter | Rat | F-TGTAGGAGCGCAGCGGAGCA<br>R-CCCACCCGTCGCCATGGAGA |
| cFOS promoter | Human | F-TGTTATAAAAGCAGTGGCTGCG<br>R-TCTTGGCTTCTCAGATGCTCG |
| cJUN promoter | Human | F-TCTCTCCGTCGCAACTTGTC<br>R-ACGCAGCAGTTGCAAACATT |
| SMARCA4 promoter | Human | F-GGGAAGTTTTGCAGAGAAGGC<br>R-CCTGGGAACCGCTTTGATCC |
